# Transcriptional elongation machinery controls vulnerability of breast cancer cells to PRC2 inhibitors

**DOI:** 10.1101/2020.01.08.898577

**Authors:** Ngai Ting Chan, Peng Liu, Junfeng Huang, Yidan Wang, Irene Ong, Lingjun Li, Wei Xu

## Abstract

CTR9 is the scaffold subunit in Paf1c, a multifunctional complex regulating multiple steps of RNA Pol II-mediated transcription. Using inducible and stable CTR9 knockdown breast cancer cell lines, we discovered that the expression of a subset of KDMs, including KDM6A and Jarid2, is strictly controlled by CTR9. Global analyses of histone modifications revealed a significant increase of H3K27me3 upon loss of CTR9. Loss of CTR9 results in a decrease of H3K4me3 and H3K36me3 in gene bodies, and elevated levels and genome-wide expansion of H3K27me3. Mechanistically, CTR9 depletion triggers a PRC2 subtype switching from PRC2.2 to PRC2.1. As a consequence, CTR9 depletion generates vulnerability that renders breast cancer cells hypersensitive to PRC2 inhibitors. Our findings that CTR9 demarcates PRC2-mediated H3K27me3 levels and genomic distribution, provide a unique mechanism of transition from transcriptionally active to repressive chromatin states and sheds light on the biological functions of CTR9 in development and cancer.

## INTRODUCTION

The Polymerase-Associated Factor 1 complex (Paf1c) was originally identified as a Pol II-interacting complex in *Saccharomyces cerevisiae* over 20 years ago. Paf1c has emerged as a highly conserved and multifunctional complex regulating multiple steps of RNA polymerase II (RNAPII)-mediated transcription (Van Oss et al., 2017). In budding yeast, Paf1c consists of five subunits, which includes Paf1, Ctr9, Cdc73, Leo1 and Rtf1. In higher eukaryotic organisms, Rtf1 is loosely attached to Paf1c, while Ski8/Wdr61 is an additional subunit of Paf1c (Zhu et al., 2005). Paf1c regulates multiple phases of transcription, including transcription elongation, transcription termination, and RNA 3’-end polyadenylation (Van Oss et al., 2017). Recent studies show that Paf1 and Ctr9 are essential for Paf1c integrity (Chu et al., 2013; Vos et al., 2018; Yu et al., 2015). Paf1c promotes RNAPII pause release. In addition, it regulates gene expression by controlling multiple transcription coupled histone modifications, including H2BK123ub, H3K4me2/3, and H3K36me2/3, through interaction with different histone modification enzymes (Tomson and Arndt, 2013). The defined functions of Paf1c in chromatin modification and transcription elongation control have marked Paf1c as an essential regulator of RNAPII transcription. Inactivation or overexpression of different subunits has been found in various cancer types, and the oncogenic or tumor suppressor function of individual subunits appears to be context-dependent (Jaehning, 2010). For example, germline mutations in *CTR9* were identified in Wilms tumor families, implicating *CTR9* as a Wilms tumor predisposition gene (Hanks et al., 2014). We have found that CTR9 is enriched in estrogen receptor α (ERα) positive breast cancers, and high expression of CTR9 correlates with poor prognosis and tamoxifen resistance (Zeng and Xu, 2015). Knocking down CTR9 in ERα+ breast cancer cells erased >90% of estrogen-regulated transcriptional response, demonstrating the function of CTR9 in promoting breast cancer progression (Zeng et al., 2016).

Polycomb repressive complex 2 (PRC2), the sole mammalian multi-subunit complex responsible for H3K27me3, is essential for maintaining cellular identity and development of multicellular organisms (Yu et al., 2019a). PRC2 is comprised of the core subunits EZH1/2, EED, SUZ12, and RbAp46/48, as well as a number of sub stoichiometric proteins (Holoch and Margueron, 2017; van Mierlo et al., 2019). Though EZH2 is the catalytic subunit of PRC2 (Wu et al., 2013), the physical interaction of EZH2 with EED and SUZ12 is necessary for full H3K27 methylation activity, as well as for modulating complex stability, and nucleosome binding ability. In the absence of EED or SUZ12, EZH2 is autoinhibited (Antonysamy et al., 2013; Cao and Zhang, 2004; Wu et al., 2013). Emerging evidence supports the existence of two different PRC2 complex subtypes in vertebrate. The subtype is determined by the auxiliary proteins associated with the core PRC2 complex (Holoch and Margueron, 2017; Yu et al., 2019a). EPOP or PALI, and one of the PCL proteins (either MTF2, PHF1, or PHF19) are found in PRC2.1, while AEBP2 and JARID2 are found in PRC2.2, (Smits et al., 2013). The overall H3K27me3 activity is deliberately balanced by PRC2.1 and PRC2.2, which are speculated to have different H3K27me3 activities (Conway et al., 2018; Oksuz et al., 2018). The auxiliary proteins are known to either promote PRC2 activity, or facilitate its recruitment to chromatin, or both. For example, JARID2, a PRC2.2-specific subunit, facilitates the recruitment of PRC2 to target genes (Pasini et al., 2010; Peng et al., 2009; Shen et al., 2009). JARID2 is also methylated by EZH2, and methylated JARID2 mimics the methylated H3 tail to stimulate PRC2 activity (Kasinath et al., 2018). The mutations on PRC2 members are frequently found in human cancers, which are often accompanied by the alteration of global levels of H3K27me2/3 (Conway et al., 2015). Elevated EZH2 levels in breast cancer are associated with poor prognosis (Gong et al., 2011; Kleer et al., 2003). Pharmacological inhibition of EZH2 is under clinical investigation for combating cancers with aberrant PRC2 activity (Kim and Roberts, 2016). Furthermore, single-cell analysis showed that loss of H3K27me3 was associated with treatment resistant breast cancer (Grosselin et al., 2019), highlighting the need to further understand how chromatin states affect drug sensitivity.

Histone lysine methylation is reversibly regulated by histone methyltransferases (HMTs), and lysine demethylases (KDMs). KDMs are classified into two distinct enzyme families (Nowak et al., 2016). KDM1A and KDM1B are members of the flavin-containing amine oxidases, whereas the rest of the KDMs with a catalytic JMJC domain belong to the 2-oxoglytarate oxygenase family (Nowak et al., 2016). KDM genes show temporal and tissue-specific expression patterns, and their expression levels and activities are stringently regulated (Lan et al., 2008). Dysregulation of KDMs, such as amplification, mutation, abnormal expression, have been implicated in breast tumorigenesis (Bamodu et al., 2016; Taube et al., 2017). For instance, aberrant expression of KDM5B and KDM6A is associated with aggressive breast cancers (Bamodu et al., 2016; Taube et al., 2017). However, the mechanisms regulating their expression remain largely unknown.

Here we identified a subset of KDMs, including PCR2 target genes KDM6A and JARID2, whose expression is precisely controlled by CTR9 levels in breast cancer cells. Global analyses of histone modifications revealed a significant increase of H3K27me3 levels upon depletion of CTR9. Loss of CTR9 induces a gradual reduction of H3K4me3 and H3K36me3 in gene bodies, followed by a drastic genome-wide increase of H3K27me3. This effect is likely attributed to switching from PRC2.2 to PRC2.1, which has high H3K27me3 activity. Moreover, exogenous expression of KDM6A or JARID2 can partially reverse the phenotypic defects in CTR9 knockdown cells. Finally, CTR9 depleted cells become addicted to H3K27me3, and are hypersensitive to PRC2 inhibition. Collectively, our study uncovers a unique mechanism by which a transcriptional elongation factor demarcates the PRC2-mediated H3K27me3 domains in breast cancer cells. The mechanism of regulation of H3K27me3 by CTR9 is likely conserved across cell types, and CTR9-dependent response to EZH2 inhibitors provides therapeutic vulnerability for breast cancer treatment.

## RESULTS

### CTR9 regulates a subset of histone lysine demethylases (KDMs) in ERα+ breast cancer cells

Data mining of the CTR9-regulated transcriptome in CTR9 inducible knockdown (KD) MCF7 cells (MCF7-tet-on-shCtr9) (Figure 1A) (Zeng and Xu, 2016) identified a subset of KDM genes that mirrors the expression of CTR9, including KDM1A, KDM2B, KDM3B, KDM5B, KDM6A and JARID2. We found that these KDM genes were down-regulated when CTR9 was knocked down by shRNA in MCF7 cells after a 7-day doxycycline (Dox) treatment (Figure 1B) regardless of whether 17β-estradiol (E2) was present or absent. RNAPII binding peaks at the transcription start site (TSS) of several KDM genes also decreased in response to CTR9 depletion (Figure S1A). We validated the transcriptome array results by RT-qPCR. Figure 1C shows that silencing of CTR9 after a 7-day Dox treatment reduced the total mRNA levels, and reduced the expression of ribosome-associated RNAs, which are indicative of actively transcribed mRNAs, of six KDMs. However, the mRNA levels of KDM4B and KDM6B did not significantly change. CTR9’s regulation of KDMs was also observed at the protein level. As shown in Figure 1D, the protein levels of six KDMs significantly decreased in the total cell lysates and extracted chromatin fractions. Consistent with their mRNA levels in response to CTR9 depletion, KDM4B and KDM6B protein levels also remained unchanged. To exclude the possibility that this observation was specific to MCF7, or due to off-target effects of anti-CTR9 shRNA, we employed MCF7 cells as well as T47D cells, another well-established ERα+ breast cancer cell line, stably expressing scramble-controlled shRNA, or two distinct lentiviral CTR9-targeted shRNAs (shCtr9#3 and #5) (Zeng and Xu, 2015). The shRNA expression led to a drastic reduction of CTR9 protein (Figure S1C-D). As a result, decreased expression at both the mRNA and protein levels was observed for the same KDMs, confirming our previous results (Figure S1B-1D).

**Figure 1.**
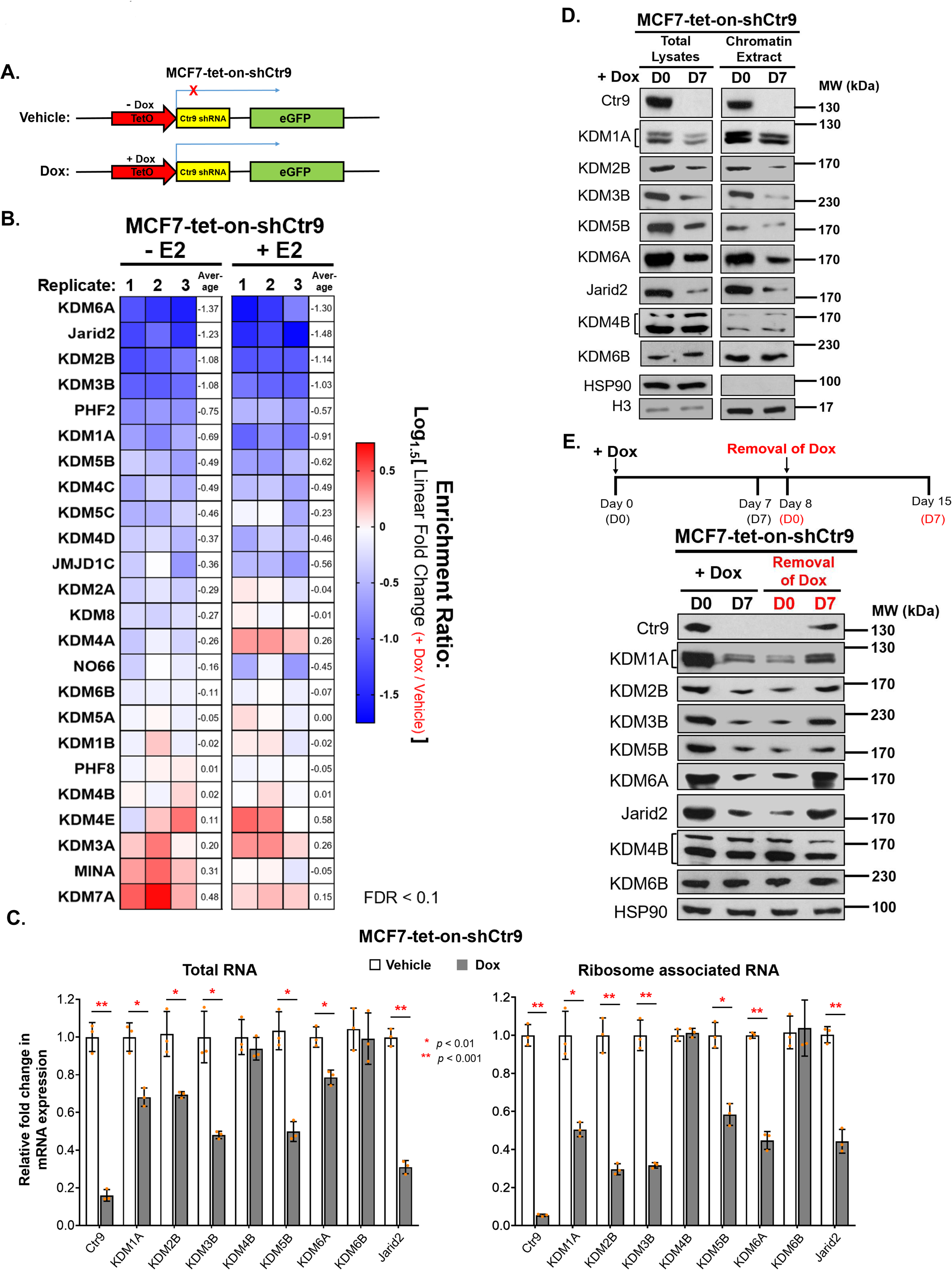
Ctr9 regulates a panel of KDM proteins in MCF7 breast cancer cells. (A) Schematic representation of the design of the doxycycline (Dox) inducible CTR9 knockdown system in MCF7 cells. eGFP reporter positive cells indicate shCTR9 RNA expression. (B) Heat map illustrating relative enrichment ratio in expression of all human histone lysine demethylases (KDM) genes measured using an Affymetrix human transcriptome array 2.0. Triplicate samples of MCF7-tet-on–shCtr9 cells were treated with vehicle or Dox in the presence of DMSO (-E2) or 10 nM E2 (+E2). (GEO accession number GSE73388) (C) RT-qPCR analysis of mRNA levels of CTR9 and KDM genes purified from either total RNA (left), or ribosome-bound RNA (right) from MCF7-tet-on–shCtr9 cells treated with either vehicle or Dox for seven days. Relative fold changes in mRNA levels are represented as the mean ± SD (n = 3). Total RNA or ribosome-bound RNAs were normalized to the internal control gene β-Actin or 18s rRNA, respectively. Values in the vehicle group were set to 1. Adjusted p-value of multiple t test (paired, two tails) with Holm-Sidak correction are calculated. (D) Western blot analysis of specific KDM proteins indicated on the left. MCF7-tet-on-shCTR9 cells were treated with 500ng/ml Dox for zero days (D0) or seven days (D7), and total cell lysates and chromatin fractions were extracted. HSP90 and histone H3 were used as loading controls for the total lysate fraction, and the chromatin fraction, respectively. (E) Western blot analysis of specific KDM proteins upon addition and withdraw of Dox (bottom) as shown in the scheme (top). MCF7-tet-on-shCtr9 cells were cultured in full medium containing 500ng/ml Dox for zero days or seven days, and then switched to medium lacking Dox (removal of Dox) and cultured for an additional seven days. HSP90 was the loading control.

To assess if decreased KDM expression in response to loss of CTR9 is reversible, we treated the MCF7-tet-on-shCtr9 cells with Dox for seven days, removed Dox on Day 8, and continued culturing the cells for seven days. CTR9 expression was gradually restored after removal of Dox, and returned to its original expression levels after seven days, as we have published previously (Zeng and Xu, 2015). The protein levels of KDMs were also found to be dynamically regulated by the addition and withdrawal of Dox, mirroring the expression pattern of CTR9. As expected, the protein levels of KDM4B and KDM6B remained constant, regardless of changes in CTR9 levels (Figure 1E). These results demonstrate that the expression of KDMs is reversibly and dynamically regulated by CTR9 in ERα+ breast cancer cell lines. To study whether the expression of CTR9 and a subset of KDMs is positively correlated in breast tumor specimens, we analyzed RNA-seq data from 593 primary ERα^+^ breast tumors in The Cancer Genome Atlas (TCGA) (Ciriello et al., 2015). We observed a statistically significant positive correlation between the expression of CTR9 and the expression of KDM1A, KDM3B, KDM5B, KDM6A and Jarid2 (Figure S1E). Collectively, these results demonstrate that CTR9 regulates a subset of KDMs in breast cancer cells.

### CTR9 determines cellular H3K27me3 levels

Since KDMs targeting different histone modification sites are regulated by CTR9, we went on to examine whether global histone modifications are altered upon CTR9 depletion. Total histones were extracted from nuclear pellets of MCF7-tet-on-shCTR9 cells after treatment with Dox for seven days, and subjected to liquid chromatography, followed by tandem mass spectrometry (LC-MS/MS). The transcription-coupled histone modifications such as H3K4me3 and H3K36me3 decreased when CTR9 was depleted as we reported previously (Zeng and Xu, 2015). Surprisingly, transcriptional repressive histone markers H3K27me2 and H3K27me3 robustly increased (Figure 2A). The histone modification changes in response to CTR9 KD were validated by western blotting (Figure 2B) where a significant increase of H3K27me3 was observed after a 7-day Dox treatment. This result suggests that loss of CTR9 results in a global elevation of H3K27me3 levels. Furthermore, the accumulation of H3K27me3 in CTR9 KD cells was not cell type specific, as this was also observed in MCF7, T47D, and BT474 cells stably expressing two distinct CTR9 targeting shRNAs (shCtr9, #3 or #5), as measured by ELISA and western blotting of purified histones (Figure S2A-C). In contrast, Dox treatment of MCF7-tet-on parental cells did not result in any changes in H3K27me1/2/3 (Figure S2D). To further interrogate whether elevation of H3K27me3 levels in CTR9 KD cells can be reversed by re-expressing CTR9 by removal of Dox, we performed a time-course Dox addition/removal treatment experiment as described above. Figure 2C shows that H3K27me3 levels increased when CTR9 was depleted, and H3K27me3 returned to its original levels when CTR9 is restored. The dynamic changes of H3K27me3 in response to CTR9 levels were further quantified by flow cytometry using an Alexa Fluor^®^ 647-labelled H3K27me3 antibody (Figure 2D) as well as quantitative mass spectrometry analyses (Figure 2E). As anticipated, H3K27me3 signal intensity increased in response to CTR9 depletion (+Dox, 7 days) as compared with untreated MCF7 cells (+ Dox, 0 days). When CTR9 was re-expressed by removing of Dox for seven days, H3K27me3 levels decreased. To visualize the H3K27me3 changes in individual cells, we performed 3D scanning of immuno-fluorescence staining of H3K27me3 (Figure 2F). The results showed that, although the response of individual cells varies, possibly due to heterogenous expression levels of shCTR9 (expression indicated by eGFP), the overall intensity of H3K27me3 staining is inversely correlated with CTR9 levels in a dynamic manner. Quantification of H3K27me3 intensity is depicted as the ratio of H3K27me3 to nucleus staining intensity from 20 selected cells with the entire nucleus in the 3D volume view (Figure 2G). Together, these data strongly suggest that CTR9 is a *bona fide* regulator of cellular H3K27me3 levels.

**Figure 2.**
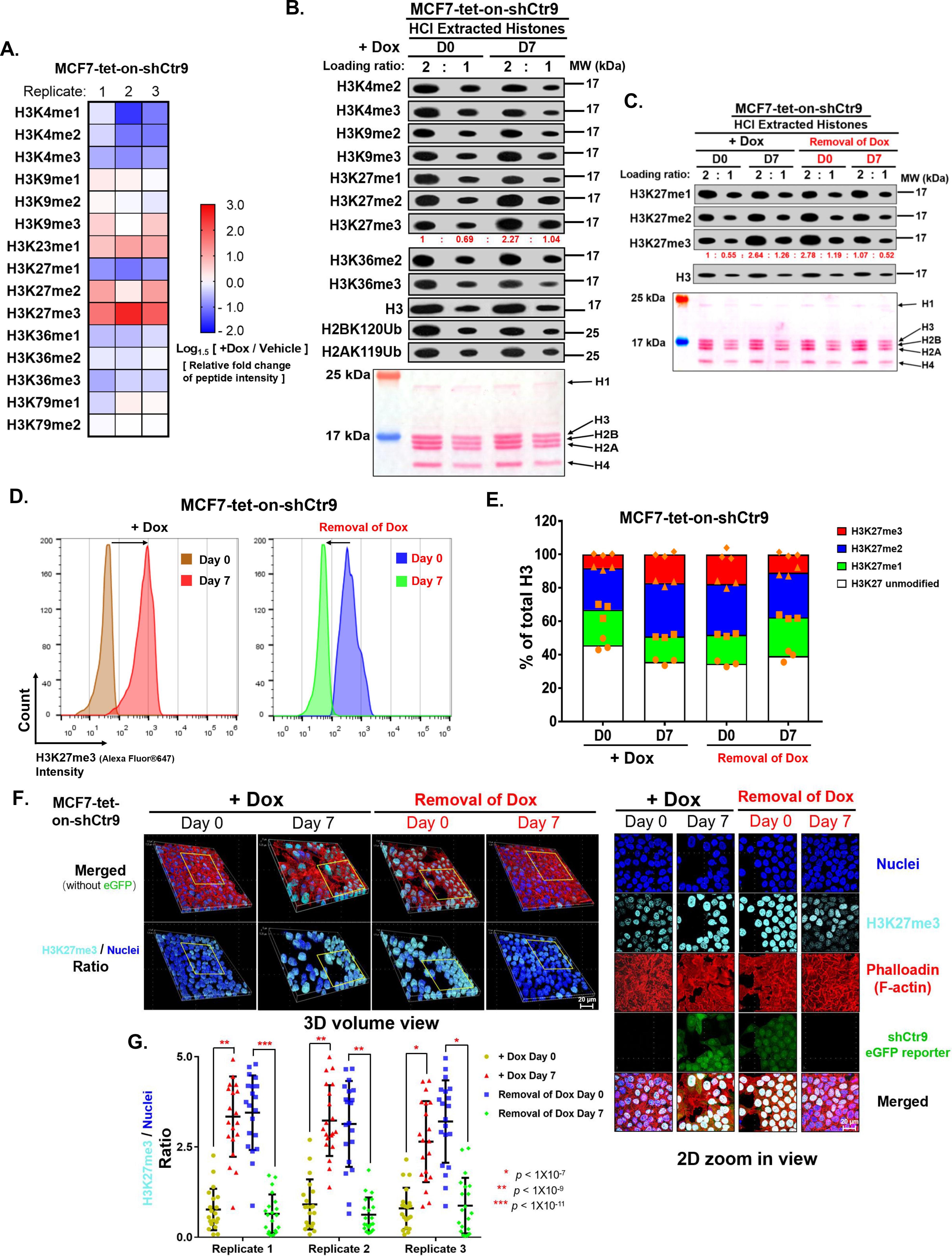
Depletion of Ctr9 leads to the global increase of H3K27me3 in MCF7. (A) Heat map showing the log2 relative fold change of histone methylation as measured by liquid chromatography tandem mass spectrometry (LC-MS/MS) in triplicate samples of MCF7-tet-on–shCtr9 cells treated with vehicle or Dox. (B) Western blot analysis of histone modifications (top). Histones were extracted by HCl from MCF7-tet-on-shCtr9 cells were treated with 500ng/ml Dox for 0 day or 7 days. Histone H3 serves as a loading control. Ponceau S staining (bottom) of HCl extracted histones MCF7-tet-on-shCtr9 cells were treated with Dox for 0 day or 7 days and loaded in two-fold dilution. The bands intensity of H3K27me3 are quantified by ImageJ after normalizing with H3 loading controls. (C) Western Blot analysis of H3K27me1/2/3 in MCF7-tet-on–shCtr9 cells treated with vehicle or Dox for 7 days followed by removal of Dox for 0 or 7 days (Top). Ponceau S staining (Bottom) of HCl extracted histones from MCF7-tet-on-shCtr9 cells in two-fold loading ratio. The bands intensity of H3K27me3 are quantified by ImageJ after normalizing with H3 loading controls. (D) Flow cytometry analysis of H3K27me3 intensity in MCF7-tet-on–shCtr9 cells upon addition and removal of Dox. (E) Bar chart showing the relative percentage of H3K27me1/2/3 as measured by liquid chromatography tandem mass spectrometry (LC-MS/MS) in triplicate samples of MCF7-tet-on–shCtr9 cells treated with vehicle or Dox for 7 days followed by removal of Dox for 0 or 7 days. (F) Representative images in 3D volume view (left) and 2D zoom view (right) of Immuno-fluorescence staining of H3K27me3/nuclei/F-actin in MCF7-tet-on– shCtr9 cells upon addition and removal of Dox. (G) Quantification of H3K27me3/nuclei staining intensity ratio in 20 randomly selected cells with complete nuclei. Data are represented as mean ± SD (n=3). Adjusted p-value of multiple t test (unpaired, two-tailed) with Holm-Sidak correction are calculated.

### CTR9 restricts genome-wide distribution of H3K27me3

PRC2 deposits H3K27me3 marks in spatially defined chromatin domains to repress gene expression. Since CTR9 KD leads to a global increase of H3K27me3 levels, we examined if CTR9 demarcates genomic H3K27me3 distribution. We performed chromatin immunoprecipitation sequencing (ChIP-seq) using antibodies against H3K27me3, H3K4me3 and H3K36me3, in CTR9 inducible and stable knockdown MCF7 cells. The use of MCF7-tet-on-shCtr9 cells allows us to monitor dynamic changes of H3K4me3, H3K36me3 and H3K27me3 in response to the gradual decrease of CTR9, while the stable CTR9 KD MCF7 cells expressing a scrambled shControl, and two distinct CTR9 shRNAs, allow us to assess permanent changes in histone modifications.

In the CTR9 inducible knockdown system, total H3K27me3 peak numbers increased over two-fold when CTR9 was silenced after a seven-day Dox treatment (Figure 3A). The H3K27me3 peaks increased across the genome and were not restricted to specific chromatin regions (i.e., promoter, exon, intron and intergenic regions) (Figure 3A). Next, we classified H3K27me3 peaks based on peak intersection, and generated four distinct clusters in the vehicle or Dox-treated groups (Figure 3B-up). Cluster I contains 7,128 H3K27me3 peaks that only exist in the vehicle-treated group. Notably, Cluster II contains a significantly higher number of H3K27me3 peaks (23,581) that are unique to the Dox treated group. Cluster III and Cluster IV represent overlapping H3K27me3 peaks where one peak in the vehicle treated group intersects with one or more peaks in the Dox treated group, or *vice versa*. Based on this peak classification, Figure 3B (bottom) depicts clustered heatmaps and line plots summarizing the average ChIP-seq signals in both vehicle and Dox-treated groups at the corresponding peak regions using the locus with the highest ChIP-seq signal as the center with ± 2kb expansion. Representative snapshots of the genome browser for each cluster are shown in Figure 3C. Because H3K27me3 peaks are too broad to define their regulated genes, we analyzed histone modification changes of 240 previously identified Ctr9 target genes in MCF7 cells. These genes were identified based on two criteria: decreased mRNA expression and reduced RNAPII binding to promoters in response to loss of CTR9 (Zeng et al., 2016; Zeng and Xu, 2015). Among the CTR9 regulated genes, 229 genes elicited either reduced H3K4me3, reduced H3K36me3 signals, or both. Only ∼10% of CTR9-regulated genes showed increased H3K27me3 signal, all of which simultaneously harbored decreased signal of either H3K4me3, H3K36me3, or both (Figure 3D, yellow and red bars). Figure 3E showed a significant reduction of H3K4me3 signal, whereas the H3K27me3 signal only modestly increased on CTR9 target genes. The negative correlation between H3K27me3 and H3K4me3/H3K36me3 across the genome was not statistically significant (data not shown), indicating that loss of active histone marks is necessary, but not sufficient, for deposition of H3K27me3 marks.

**Figure 3.**
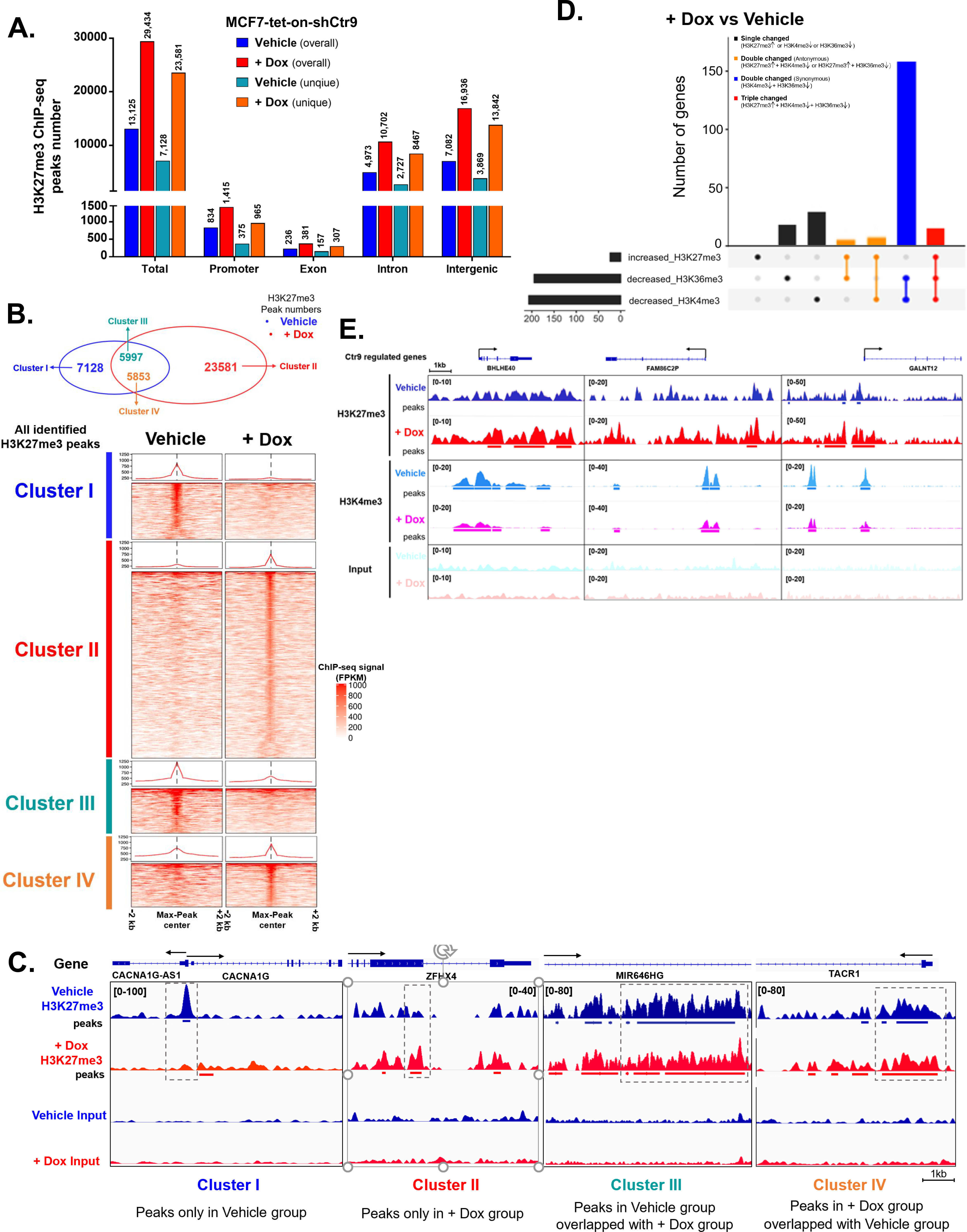
Inducible knockdown of Ctr9 results in gradual loss of H3K4me3 and H3K36me3 followed by genome-wide gain of H3K27me3. (A) Summary of overall and unique H3K27me3 peak numbers in MCF7-tet-on-shCtr9 cells treated with vehicle or Dox. Peak numbers are classified in total or in Promoter or Exon or Intron or Intergenic regions. (B) Venn Diagram showing the overlapped and unique H3K27me3 peaks between Vehicle and + Dox groups (Top). Different clusters were assigned according to the Venn Diagram intersection. Clustered heatmap of H3K27me3 ChIP-seq signal in both vehicle and + Dox groups using the maximum peak value as matrix-center (Bottom). Each cluster have a line plot on the top summarizing the average H3K27me3 ChIP-seq signal in the range of max peak center ± 2kb. (C) Representative genome browser snapshots of the H3K27me3 ChIP-seq signals and corresponding peak profile in each cluster. (D) Summary of gene numbers with increased H3K27me3 ChIP-seq signal or/and decreased H3K4/K36me3 ChIP-seq signal after inducible loss of Ctr9 (+Dox vs vehicle) within previously identified 240 Ctr9 regulated genes. (E) Representative genome-browser snapshots of serval Ctr9 regulated genes with significant increase of H3K27me3 ChIP-seq signal and decrease of H3K4me3 ChIP-seq signal simultaneously when Dox treated group was compared to vehicle.

In comparison to the inducible CTR9 KD cells, the global increase of H3K27me3 peak numbers was much more pronounced in CTR9 stable KD MCF7 cells (shCtr9#3 and shCtr9#5) when peaks were normalized to control shRNA expressing cells (Figure S3A). The increased H3K27me3 peaks spread across the genome (Figure S3A) is indicative of expansion of H3K27me3 domains. In comparison with the CTR9-inducible system, Cluster I H3K27me3 peaks decreased from 7,128 to 3,935 and 2,787 in the shControl cells, respectively (Figure 4A). In contrast, Cluster II H3K27me3 peaks increased from 23,581 to 40,595 and 53,078, respectively, for shCtr9#3 and shCtr9#5 (Figure 4B). In addition, the peak intensity of H3K27me3 within those TSSs ± 5kb regions with gained H3K27me3 peaks after CTR KD also significantly increased in CTR9 stable KD cells (Figure S3B), as well as the H3K27me3 peak width across the genome in intergenic, exon, intron, and promoter regions (Figure S3C). The five-fold increase of total H3K27me3 peaks, increased peak intensities, and broadened peak width in response to permanent CTR9 knockdown indicate that loss of CTR9 resulted in significant expansion of H3K27me3 domains in chromatin. In contrast to what was found in the inducible CTR9 KD cells, a statistically significant, negative correlation between H3K27me3 and H3K4me3 / H3K36me3 was detected at the genome-wide level in CTR9 stable KD cells (Figure S3D). With regard to 240 CTR9-regulated genes, significant number of genes (∼30 for shCtr9#3 and ∼80 for shCtr9#5) showed increased H3K27me3 signals in stable CTR9 KD cells (Figure 4B), and the increase of H3K27me3 is often coupled with decreased H3K36me3 but less prominent with decreased H3K4me3 (Figure 4B). Previous studies have shown that H3K27me3 and H3K36me2 could co-localize in some genomic regions, and depletion of H3K36me2 is accompanied by deposition of H3K27me3 (Brien et al., 2012; Streubel et al., 2018). We hypothesized that upon loss of CTR9, H3K4me3 and H3K36me3 in gene bodies decreases followed by deposition of H3K27me3 marks, and the expansion of H3K27me3 domains.

**Figure 4.**
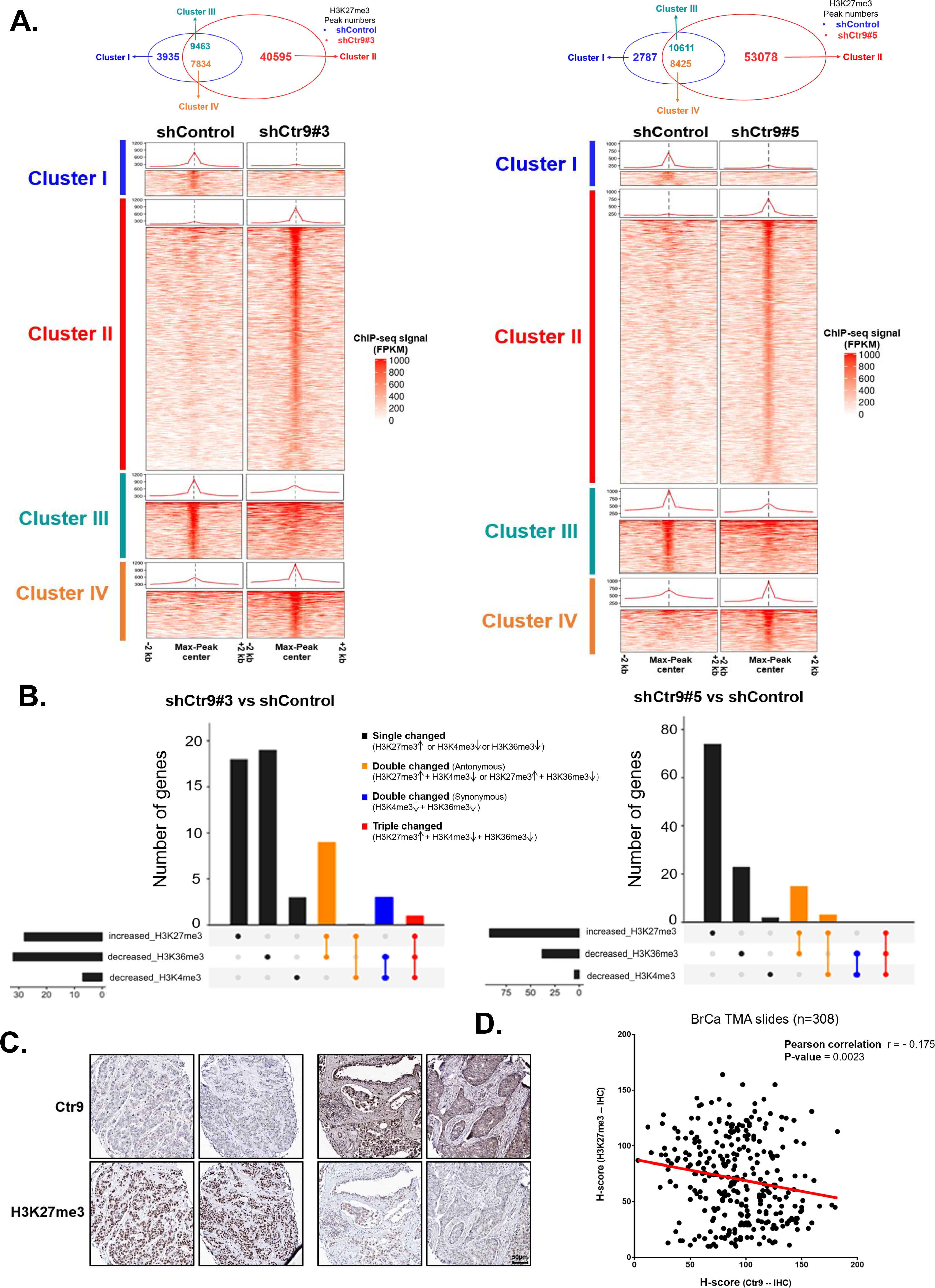
Stable depletion of Ctr9 results in drastic expansion of H3K27me3 peaks across genome. (A) Venn Diagram showing the overlapped and unique H3K27me3 peaks between Control KD and two Ctr9 KD (shRNA#3 or shRNA #5) groups (Up). Different clusters were assigned according to Venn Diagram intersection. Clustered heatmap of H3K27me3 ChIP-seq signal in both Control KD and two Ctr9 KD groups using the maximum peak value as matrix-center (Bottom). Each cluster have a line plot on the top summarizing the average H3K27me3 ChIP-seq signal in the range of max peak center ± 2kb. (B) Summary of gene numbers with increased H3K27me3 ChIP-seq signal or/and decreased H3K4/K36me3 ChIP-seq signal after stable knockdown of Ctr9. (shCtr9#3 or shCtr9#5 vs shControl) within 240 Ctr9 regulated genes (C) Representative images of 40x scanned tumor cores with high or low Ctr9 IHC staining and corresponding H3K27me3 IHC staining in breast cancer TMA slides. (D) Correlation analysis between Ctr9 and H3K27me3 IHC staining by calculating the 4-bin H-score after excluding non-available samples.(n=308) (0^+^: negative staining; 1^+^: weak staining; 2^+^: medium staining; 3^+^: strong staining). Pearson correlation r value and two-tailed t test p-value were shown.

We further investigate the correlation between CTR9 and H3K27me3 level in breast tumors using human breast cancer tissue microarrays (TMAs) containing over 300 breast tumor specimens. After optimizing staining condition with antibodies to CTR9 and H3K27me3 (Figure S4A), the intensity of immunohistochemistry staining (IHC) was determined by H-score graded optical density in the nucleus of epithelial cells in specimen (Figure S4B). Representative examples of breast tumor cores with high or low CTR9 IHC staining and corresponding inverse H3K27me3 IHC staining were shown in Figure 4C. The H-score based correlation analysis revealed a significant reverse correlation between CTR9 expression and H3K27me3 abundance (Figure 4D).

We next investigated whether CTR9-regulated KDM genes showed dynamic histone modification changes. Indeed, KDM genes showed decreased H3K4me3 peaks near gene promoters (Figure S5A) when CTR9 was inducibly knocked down by Dox-treatment and H3K27me3 peaks increased (Figure S5B). In contrast, KDM4B, a non-CTR9 target gene, did not show significant changes in H3K4me3 or H3K27me3 enrichment.

### CTR9 level is a determinant of PRC2 complex subtype

Because H3K27me3 is mainly established by the PRC2-EZH2 complex, we examined if PRC2-EZH2 repressed genes in breast cancer cells also gain H3K27me3 signal, as observed for CTR9 target genes. A recent study identified a panel of EZH2 repressed genes in a mouse breast cancer model, and *FOXC1* was shown to be repressed by H3K27me3 in an EZH2-dependent manner (Hirukawa et al., 2018). Analyses of H3K27me3 ChIP-seq signal in the human analogues of the mouse genes showed that approximately half of the PRC2-repressed genes displayed a dramatic increase of H3K27me3 in CTR9 KD cells (Figure S6A). A snapshot of the genome browser for H3K27me3 signals at *FOXC1* in shControl and shCTR9 stable KD cells illustrated a dramatic increase of H3K27me3 in *FOXC1* upon loss of CTR9 (Figure S6B). These results demonstrate that CTR9 regulates PRC2 target genes in a H3K27me3-dependent manner.

The increased total H3K27me3 levels, and broadened H3K27me3 peak-width (Figure 4A and Figure S3C) in CTR9 KD cells conforms with enhanced PRC2 activity. The PRC2.1 and PRC2.2 subtypes were shown to antagonize each other. For instance, loss of PRC2.2 specific subunit AEBP2 led to an increase in the amount of PALI1-containing PRC2.1 (Conway et al., 2018). Moreover, H3K27me3 is elevated in AEBP2 KO ESCs (Ciferri et al., 2012). Because the levels of JARID2, another PRC2.2-specific subunit, decreased significantly in CTR9 KD cells, we speculate that CTR9 KD may cause PRC2 subtype switching by shifting the composition of PRC2 facultative components. Indeed, depleting CTR9 in MCF7 cells resulted in a dramatic increase in PRC2.1 facultative subunits PCL2, PCL3, and EPOP, and a decrease in PRC2.2 facultative subunits JARID2 and AEBP2 (Figure 5A) in total cell lysates and chromatin fractions, whereas the levels of four core subunits remained unchanged. Similar observations were made in two stable Ctr9 KD MCF7 cell lines (Figure 5B). To investigate if regulation of PRC facultative subunits by Ctr9 is reversible, we measured expression of five facultative subunits of respective PRC2 subtypes in MCF7-tet-on-shCtr9 cells. The expression of PRC2 facultative subunits, and CTR9 protein levels was strongly correlated (Figure 5C). The decrease of PRC2.2-specific subunits JARID2 and AEBP2 could be partially restored by CTR9 re-expression, accompanied by the opposite changes in PRC2.1-specific PCL2, PCL3 and EPOP proteins. The negative correlation between CTR9 and PRC2.1 facultative subunits, PCL1/3, and EPOP, at the mRNA level was observed in 803 ER+ primary breast tumor samples in TCGA (Figure 5D). To test if the inverted levels of PRC2 facultative subunits in response to changes in CTR9 levels resulted in switching of PRC2 subtypes, we performed co-immunoprecipitation using antibodies against the core subunits of the PRC2 complex EZH2 and SUZ12 to pull down other core and auxiliary proteins from nuclear extract. Figure 5E shows that, in the presence of CTR9, PRC2 core subunits tended to precipitate PRC2.2-specific JARID2 and AEBP2. When CTR9 was depleted, PRC2.1-specific subunits PCL2/3 and EPOP were enriched instead. We reason that this switch is likely due to the relative abundance of facultative subunits corresponding to CTR9 levels. In addition, the inversely correlated expression levels of CTR9 and H3K27me3 levels can also be detected in breast cancer cell lines (Figure 5F). CTR9 levels are higher in ERα^+^ than triple-negative breast cancer cell lines (TNBC) as we published previously (Zeng and Xu, 2015), whereas H3K27me3 levels are higher in TNBC. In summary, CTR9 expression is a key determinant for PRC2 complex composition and H3K27me3 activity.

**Figure 5.**
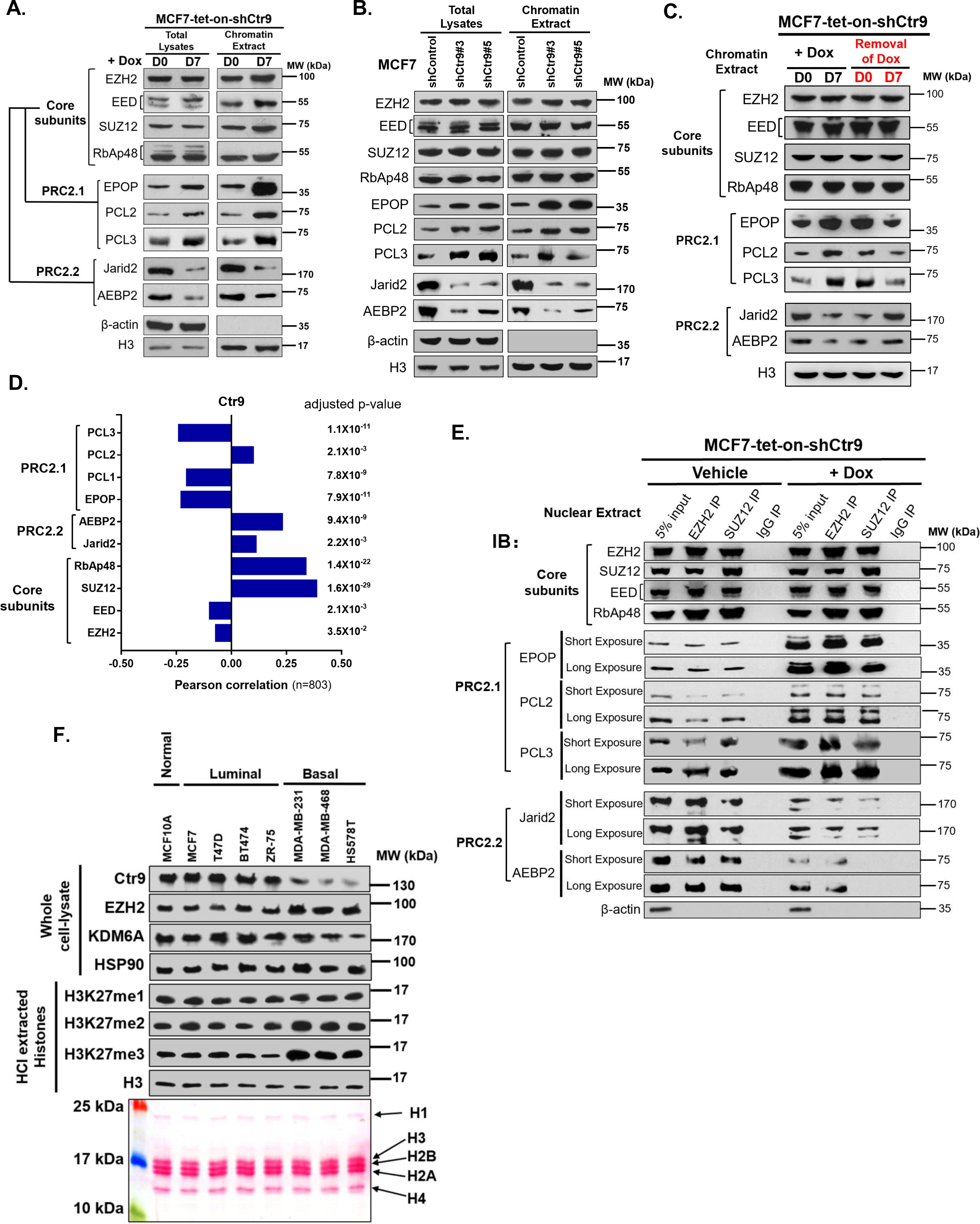
Depletion of Ctr9 results in switching from PRC2.2 to PRC2.1 subtype. (A) Western Blot analysis of PRC2 core (EZH2/SUZ12/EED/RbAp48) and facultative subunits (PRC2.1: EPOP/PCL2/PCL3; PRC2.2: Jarid2/AEBP2) in total lysates and chromatin fractions in MCF7-tet-on-shCtr9 cells treated with 500ng/ml Dox for zero days (D0) or seven days (D7). β-actin and Histone H3 were used as loading controls for total lysates and chromatin extract, respectively. (B) Western blotting analysis of PRC2 core and facultative subunits in total lysates and extracted chromatin fraction of MCF7-shControl or MCF7-shCtr9#3 or ShCtr9#5 cells. Histone H3 was used as a loading control. (C) Western Blot analysis of PRC2 core subunits and facultative components using chromatin extracts from MCF7-tet-on-shCtr9 cells were addition and withdraw of Dox. Histone H3 was used as a loading control. (D) Bar chart showing the person correlation of RNA-seq data between Ctr9 and core or facultative subunits of PRC2 complex in 803 primary ER^+^ breast tumor samples from TCGA. P-values were adjusted by Benjamini & Hochberg correction. (E) Co-Immunoprecipitation of PRC2 core subunits (EZH2/SUZ12/EED/RbAp48) and auxiliary components (EPOP/PCL2/PCL3; Jarid2/AEBP2) using nuclear extracts from MCF7-tet-on-shCtr9 treated with vehicle or Dox for 7 days. IgG IP served as negative controls. β-actin was used as a loading control for input. (F) Western blot of Ctr9, EZH2, KDM6A and H3K27me1/2/3 in an immortalized non-transformed mammary epithelial cell line (MCF10A), four ER^+^ human breast cancer cell lines, and three TNBC cell lines. HSP90 and histone H3 were loading controls for whole cell lysate and extracted histones, respectively.

### Both JARID2 and KDM6A mediate the biological activities of CTR9

Among the six KDMs regulated by CTR9, JARID2 and KDM6A are both H3K27me3 modulators. JARID2 is part of the H3K27me3 ‘writer’ complex PRC2 (Peng et al., 2009) and KDM6A is an H3K27me3 ‘eraser’ (Agger et al., 2007). As CTR9 KD in MCF7 cells resulted in a number of growth-related phenotypes, including cell cycle arrest, reduced colony formation, and senescence (Zeng and Xu, 2015), we tested if exogenous expression of either JARID2 or KDM6A (+Flag-Jarid2/KDM6A) in CTR9 KD cells (Figure 6A) could rescue these phenotypes, where re-expression of Ctr9 (+ Flag-Ctr9) serves as a positive control. Our results showed that exogenous expression of either JARID2 or KDM6A partially rescued the defects in growth (Figure 6B) and colony formation (Figure 6C) caused by CTR9 knockdown. Consistent with our published work (Zeng and Xu, 2015), CTR9 KD MCF7 cells proliferated slower, as indicated by a decreased population of cells in S phase of the cell cycle. Both effects could be partially rescued by exogenously expressing JARID2 or KDM6A, although not as effective as Ctr9 restoration, as observed by cell cycle analyses using propidium iodide (PI) (Figure 6D) and Edu staining (Figure S7A). Importantly, exogenous expression of either JARID2 or KDM6A partially reduced the elevated H3K27me3 level caused by loss of Ctr9, where re-expression of Ctr9 serves as a positive control.

**Figure 6.**
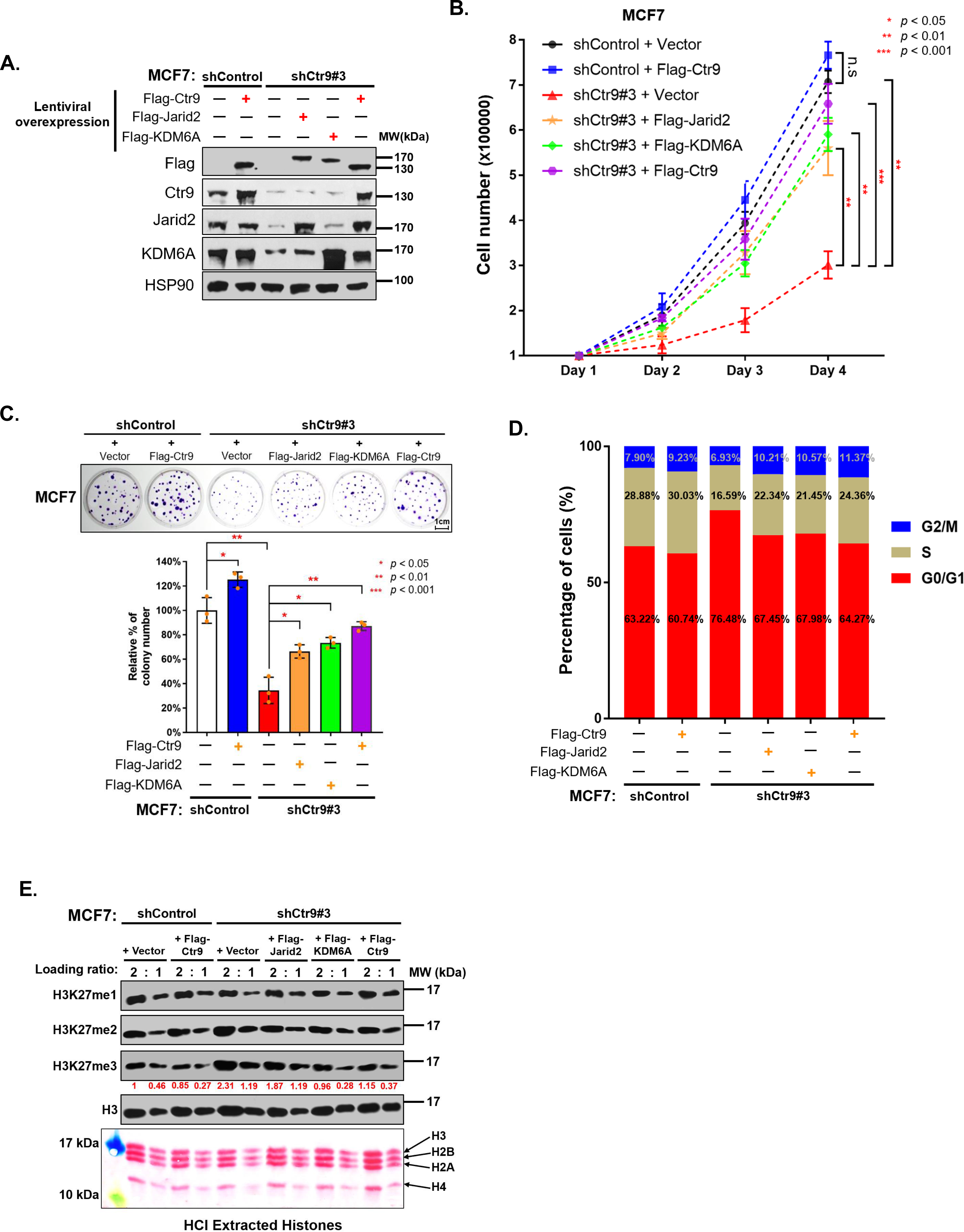
Exogenous expression of JARID2 or KDM6A partially rescues the cellular defects caused by Ctr9 KD. (A) Western Blot validation of exogenous expression of Flag-tagged Ctr9, Jarid2 or KDM6A in MCF7-shControl or MCF7-shCtr9#3 cells. MCF7-shControl or MCF7-shCtr9#3 cells transfected with blank vector served as negative controls. β-actin was used as a loading control. (B) Measurement of cell proliferation by cell counting in Control KD or Ctr9 KD MCF7 cells transfected with blank vector or Flag-Jarid2/KDM6A/Ctr9 expressing plasmid. Adjusted p-value of multiple t test (unpaired, two-tailed) with Holm-Sidak correction are calculated. (C) Representative image of 2D colony formation of Control KD or Ctr9 KD MCF7 cells transfected with blank vector or Jarid2/KDM6A overexpression plasmid (n=3) (Top). Quantification of relative colony numbers using Image J normalized by average colony diameters (Bottom). Number in MCF7-shControl transfected with blank vector group was set as 100%. Data are represented as mean±SD (n=3). P-value of unpaired two-tailed t test with Welch correction are shown. (D)PI staining-based cell cycle analysis of control KD or Ctr9 KD MCF7 cells transfected with blank vector or Flag-Jarid2/KDM6A/Ctr9 expressing plasmid. (E) Western Blot of H3K27me1/2/3 in MCF7-shControl or MCF7-shCtr9#3 cells with exogenous expression of Flag-tagged Ctr9, Jarid2 or KDM6A (top). Ponceau S staining of HCl extracted histones in two-fold dilution for loading (bottom). The bands intensity of H3K27me3 are quantified by ImageJ after normalizing with H3 loading controls.

All data support that CTR9 has a profound impact on H3K27me3 by regulating both ‘writer’ and ‘eraser’ enzymes, and that JARID2 and KDM6A are downstream effectors mediating some CTR9 cellular functions in breast cancer cells.

### CTR9 depletion sensitizes breast cancer cells to PRC2 complex inhibitors

High levels of EZH2 and H3K27me3 in ER^-^ breast cancer patients predict poor overall survival, and EZH2 inhibitor GSK343 has been shown eliciting robust inhibition of ER^-^ breast tumor growth in preclinical model (Yu et al., 2019b). The remarkable increase of total H3K27me3 levels upon depletion of CTR9 implies an addiction of cells to H3K27me3 for survival. If this were true, we expect that CTR9 knockdown cells would elicit higher sensitivity to EZH2 inhibitors than parental cells. Cell viability was measured after treatment with UNC1999, a chemical inhibitor targeting both EZH2 and EZH1, and UNC2400, a structurally similar but inactive analog compound, to exclude the possibility of off-target effects (Konze et al., 2013). As expected, CTR9 KD MCF7 cells were more sensitive to UNC1999 than parental MCF7 cells, as shown by MTT assays, whereas EZH2 KD MCF7 cells (shEZH2) were insensitive to UNC1999, serving as a negative control (Figure 7A). A similar result was observed by cell counting (Figure 7B) and colony formation assays (Figure 7C). UNC1999 inhibited the growth of CTR9 KD MCF7 cells in a dose-dependent manner, whereas the negative analog UNC2400 showed no cytotoxic effects. To exclude a drug-specific effect, we tested two additional mechanistically distinct PRC2 inhibitors, GSK343 and EED 226 (Qi et al., 2017; Verma et al., 2012). The results (Figure S8A-B) were similar to those of UNC1999. Collectively, our data indicate that depletion of CTR9 leads to increased sensitivity towards the PRC2 inhibitors. To determine whether the EZH2 inhibitor causes elevated apoptosis or necrosis upon CTR9 depletion, we performed flow cytometry analyses after labeling cells with PI and annexin V-FITC (Figure 7D). The dose-responsive increase of apoptosis by UNC1999 was quantified in Figure 7E. Both apoptotic and necrotic cells were detected in CTR9 KD cells, but not in the shControl cells after 5 μM UNC1999 treatment. When the concentration of UNC1999 increased to 12.5 μM, nearly all of the Ctr9 KD cells became apoptotic or necrotic; however, ∼30% of control KD cells remained viable (Figure 7E). The higher sensitivity against EZH2 inhibitors when loss of CTR9 was not only observed in 2D monolayer but also in 3D tumor spheroids. When treated with ascending concentration of UNC1999 from 1 μM to 50 μM for 2 days, CTR9 KD MCF7 3D spheroids (shCtr9#3/shCtr9#5) elicit stronger cytotoxic response (more red cells and less green cells) than control knockdown group (shControl). UNC2400 (50 μM) serves as a negative control (Figure S8C). A time dependent measurement of 3D spheroids-area were recorded when treating with high dose of UNC1999 (50 μM) for 0 – 72 hours (Figure S8D). The results shown that the CTR9 depleted spheroids (shCtr9#3/shCtr9#5) disassemble in a much rapid manner in contrast to the control group (shControl), whereas 50 μM UNC2400 treatment does not affect the integrity of 3D spheroids for both CTR9 KD and control KD groups. Collectively, EZH2 inhibitor UNC1999 can inhibit growth and induce apoptosis in MCF7 cells and increase their sensitivity to EZH2 inhibitors depends on the levels of CTR9. CTR9-depleted cells gain H3K27me3 and elicit stronger sensitivity to EZH2 inhibitors than parental cells in both 2D monolayer and 3D spheroid models.

**Figure 7.**
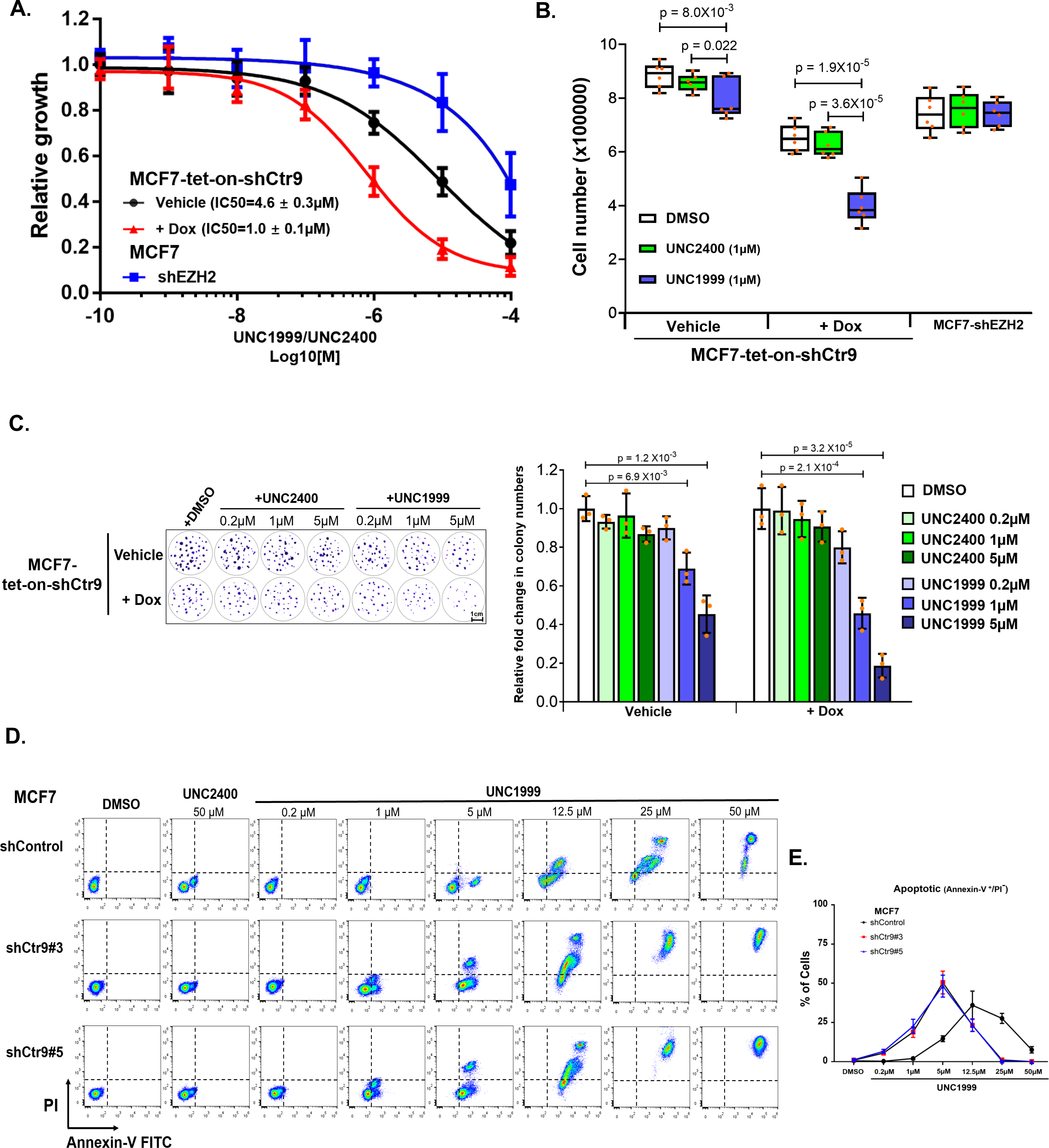
Depletion of Ctr9 sensitizes MCF7 cells to UNC1999, an EZH2 inhibitor. (A) Cell viability measured by MTT assays after treating MCF7-shEZH2 or MCF7-tet-on-shCtr9 cells (vehicle or Dox) with increasing doses of UNC1999. Data are represented as mean±SD (n=3) normalized to a control compound UNC2400 at corresponding concentrations. IC_50_ was calculated through non-linear regression against log(inhibitor) response model. (B) Measurement of cell proliferation by cell counting with trypan blue exclusion after treating MCF7-shEZH2 or MCF7-tet-on-shCtr9 cells (vehicle or Dox) with DMSO control, 1 μM UNC2400 or 1μM UNC1999 for 4 days. Data are represented as mean±SD (n=6). Adjusted p-value of multiple t test (unpaired, two-tailed) with Holm-Sidak correction are shown. (C) Representative image of 2D colony formation of MCF7-tet-on-shCtr9 cells (vehicle or Dox) treated with DMSO or 1 to 10μM of UNC2400/UNC1999 for 2 weeks (n=3) (Left). Relative colony numbers were quantified using Image J normalized by average colony diameters (Right). DMSO treatment groups were set as 100%. Data are represented as mean±SD (n=3). Adjusted p-value of multiple t test (unpaired, two-tailed) with Holm-Sidak correction are shown. (D) Flow cytometry analyses of PI uptake and annexin-V FITC labeling in MCF7-shControl or MCF7-shCtr9#3 or Ctr9#5 cells treated with DMSO, 0.2 to 50 μM UNC1999, or 50 μM UNC2400 for 2 days. (E) Flow cytometric quantification of percentage of apoptotic cells (Annexin-V^+^/PI^-^) in MCF7-shControl or MCF7-shCtr9#3 or MCF7shCtr9#5 after treating with DMSO or ascending concentrations of UNC1999.

## DISCUSSION

Here we report that CTR9 governs the establishment of H3K27me3 repressive domains, beyond its well characterized functions in transcriptional regulation and transcription-coupled histone modifications (i.e., H2Bub, H3K4me3, H3K36me3). This discovery provides an explanation for the discrepancy between the phenotypes observed in lower eukaryotes (i.e. yeast) and multicellular organisms when CTR9 is depleted. While Ctr9 is not an essential gene in *S. cerevisiae* (Chen et al., 2002; Massoni-Laporte et al., 2012), CTR9 knockout causes early embryonic lethality in higher eukaryotic organisms such as *Drosophila* (Chaturvedi et al., 2016), zebrafish (Akanuma et al., 2007) and mouse (Zhang et al., 2013). The requirement of CTR9 in preimplantation development of mice resembles the phenotypes that result from lacking core PRC2 subunits SUZ12, EZH2 and EED (Pasini et al., 2004). While PRC2 and H3K27me3 play crucial roles for establishing repressive chromatin regions, and safeguarding cell identity in multicellular organisms (Holoch and Margueron, 2017), neither PRC2, nor histone H3K27me3, exists in yeast. Our results indicate that while CTR9 maintains transcriptional control across species, controlling PRC-repressive domains is a new function acquired over the course of evolution. We surmise that the embryonic lethality of CTR9-null metazoans is more likely attributed to deregulated H3K27me3 than transcription inhibition *per se*.

Among the KDM family proteins, expression of six KDMs (KDM1A/LSD1, KDM2B, KDM3B, KDM5B, KDM6A and Jarid2) was regulated by Ctr9. KDM4B and KDM6B, on the other hand, are not Ctr9 target genes. The regulation of KDMs by Ctr9 is reversible as shown in the Ctr9 shRNA inducible system where the levels of KDMs decreased upon Ctr9 was silenced and increased upon Ctr9 was restored (Figure 1E). Moreover, the regulation of KDMs by Ctr9 is not an off-target effect of shRNA, neither it is cell-line specific (Figure S1B-D). At first glance, the Ctr9-regulated KDMs do not seem to share common properties. These KDMs span two enzyme families, are not under the same phylogenetic trees, and have distinct histone lysine modification substrates. Further analyses reveal that Ctr9-regulated KDMs are PRC2 targets, since the repression and activation of these KDMs are accompanied by the decrease of H3K4me3 and increase of H3K27me3 (Figure S4) in coding regions. In contrast, H3K27me3 was not detected on non-PRC regulated KDMs such as KDM4B. Because the KDMs have diverse histone and non-histone substrates, the identification of a subset of KDMs as PRC2 target genes and subjected to Ctr9 regulation indicates the long-term epigenetic effects on gene expression and chromatin landscape imposed by Ctr9. Among the Ctr9-regulated KDMs, KDM1A/LSD1, KDM2B/FBXL10, KDM5B/Jarid1B are overexpressed in cancers (Kooistra et al., 2012). Correspondingly, chemical inhibitors are under active development to intervene their enzymatic activities for cancer therapy. For instance, the KDM1A inhibitors are in Phase I study for treatment of acute leukemia and small cell lung cancer (Maes et al., 2015). Understanding the regulatory mechanism of KDMs would provide a better grip on the biological role of KDMs and improve pharmacological inhibition of their activities in disease states.

CTR9 is not among the 148 PRC2 interaction partners that are enriched in EZH2 and SUZ12 immunoprecipitants in ESCs (Streubel et al., 2018). Therefore, the significantly elevated levels and genome-wide distribution of H3K27me3 upon loss of CTR9 is unexpected, which implies that, despite no physical interaction, CTR9 is functionally related to PRC2. Given that CTR9 is the scaffold protein of Paf1c that regulates multiple steps of transcription, and that a decrease in KDM6A and JARID2 levels is accompanied by loss of CTR9, we envision that at least three mechanisms may collectively contribute to the establishment of H3K27me3 domains: inhibition of transcription, decreased H3K27me3 demethylation, and alteration of PRC2 activity.

First, inhibition of transcription alone has been shown to be sufficient for gain of H3K27me3 in the gene body (Hosogane et al., 2016; Riising et al., 2014). In mouse ESCs, transcriptional inhibition by RNAPII inhibitors 5,6-dichloro-1-beta-D-ribofurano-sylbenzimidazole riboside (DRB), and triptolide, induces genome-wide ectopic PRC2 recruitment (Riising et al., 2014). Hosogane M. *et al*. also reported that abrogation of transcription induces accumulation of H3K27me3 in the gene body (Hosogane et al., 2016). We have previously shown that RNAPII binding, H3K4me3, and H3K36me3, are drastically decreased by knocking down CTR9 in MCF7 cells (Zeng et al., 2016). Comparing the histone modification changes in MCF7 CTR9-inducible and stable CTR9 knockdown systems, we observed a stepwise loss of H3K4me3, and H3K36me3, as well as a gain of H3K27me3 signals on CTR9 regulated genes (Figure 3D and 4B), reinforcing that depletion of activating histone marks precedes the deposition of H3K27me3. This is also consistent with the role of the active histone marks in inhibiting PRC2 activity in transcribed regions (Laugesen et al., 2019) and that both H3K4me3 and H3K36me2/3 are inhibitory to PRC2 activity (Ballare et al., 2012; Schmitges et al., 2011). Because H3K4me3 and H3K36me3 are associated with actively transcribed genes, transcription inhibition by loss of CTR9 could result in H3K27me3 accumulation in gene bodies.

Second, H3K27me3 levels are dynamically regulated by the “writer” PRC2 complex, and the “eraser” KDM6 family of histone demethylases. The KDM6 family of H3K27me3 demethylases includes KDM6A (UTX), KDM6B (JMJD3), and UTY (Kooistra and Helin, 2012). CTR9 regulates the expression of KDM6A and JARID2 (Figure 1). It is unlikely that decreasing KDM6A expression alone results in an increase of H3K27me3 levels, because knocking out both KDM6A and KDM6B does not lead to a global increase in H3K27me3 levels. Instead, a small set of genes showed accumulation of H3K27me3 during development (Shpargel et al., 2014). Similarly, in T-cell acute lymphoblastic leukemia (T-ALL), KDM6A/UTX deficiency does not alter the global level of H3K27me3, but instead, causes a genome-wide redistribution of H3K27me3 (Ntziachristos et al., 2014). Only KDM6A, but not KDM6B, is regulated by CTR9. Thus, decreased KDM6A in CTR9 KD MCF7 cells may partially contribute to H3K27me3 genomic distribution, but is unlikely to be the cause of the global increase of H3K27me3 levels observed.

Third, we envision that the PRC2 subtype switching caused by decreased JARID2 in response to CTR9 depletion is the major mechanism for the global H3K27me3 increase observed. JARID2 is a distinguished member of the JMJC family of KDMs because it has no enzymatic activity of its own. Rather, it is a PRC2.2-specific subunit. A previous study showed that depletion of AEBP2 results in up-regulation of PALI1/2 (Conway et al., 2018), and conversion from JARID2-containing PRC2.2, to PCL2-containing PRC2.1 complex (Grijzenhout et al., 2016). Moreover, JARID2 knockout in differentiated ESCs causes aberrant deposition of H3K27me3 in intergenic regions of the genome (Sanulli et al., 2015). This is analogous to the effects of loss of CTR9, where JARID2 and AEBP2 levels decrease, while H3K27me3 domains in intergenic regions increase (Figures 3B and 4A), and PRC2 subtype switching from PRC2.2 to PRC2.1 (Figure 5). Therefore, we reason that the loss of CTR9 resulted in a decrease of JARID2, leading to formation of the more active PRC2.1 complex, and elevated H3K27me3 levels. Finally, exogenous expression of either KDM6A or JARID2 partially restores the cellular phenotypes in CTR9 knockdown MCF7 cells, suggesting that both KDM6A and JARID2 are the downstream effectors of CTR9, and mediate the cellular effects of CTR9 in breast cancer cells. Collectively, our studies reveal complex mechanisms by which CTR9 demarcates PRC2-mediated H3K27me3 domains, and the previously unidentified interplay between transcriptional activation machinery and transcriptional repressive complexes. Although these findings were made in breast cancer cell lines, the general principal for the establishment of H3K27me3 domains governed by CTR9 is likely conserved in other biological systems (e.g., ESCs).

Mutations in subunits of PRC2 have been increasingly identified in multiple cancers, resulting in changes in the global levels, as well as the genome-wide distribution of H3K27me3 (Conway et al., 2015). These changes in cancer cells often confer context dependent blocks to cellular differentiation and promote oncogenic signaling pathways. Either overexpression of EZH2, or inactivation of negative regulators of PRC2, augments the dependency of cancer cells on H3K27me3 and PRC2, indicating that PRC2 inhibition can increase cancer cell therapeutic vulnerability (Xu et al., 2015). For example, acute myeloid leukemia and multiple myeloma cell lines with KDM6A mutations are more sensitive to EZH2 inhibitors than KDM6A wild-type expressing lines (Ezponda et al., 2017; Van der Meulen et al., 2015). We found that either inducible or permanent CTR9 KD MCF7 cells elicit increased sensitivity to PRC2 inhibition, indicating that loss of CTR9 renders cells more addicted to H3K27me3 and PRC2. GSK126 and EPZ-6438, two highly specific EZH2 inhibitors, are currently in clinical investigation for treating lymphomas (Kim and Roberts, 2016). Because depletion of EZH2 suppressed ER-negative tumor growth and metastasis in preclinical models (Moore et al., 2013), EZH2 has emerged as a potential therapeutic target for TNBC. Our data that TNBC cell lines display higher levels of EZH2 and H3K27me3 (Figure 5F) support the application of PRC2 inhibitors in treating TNBC. Moreover, depletion of CTR9 in ER^+^ cells increases sensitivity of cells to PRC2 inhibition by >10 fold, suggesting that EZH2 inhibitors may also be applicable to ER-positive breast cancer with lower levels of CTR9. We speculate that CTR9 levels, rather than ER status, is a predictive biomarker for PRC2 dependency in breast cancer cells. Since Ctr9 depletion generates therapeutic vulnerability to pharmacological inhibition of PRC2, the CTR9 expression levels may be used as a guideline for predicting PRC2 dependency and EZH2 inhibitor sensitivity in broad cancer types.

Our findings that CTR9 demarcates PRC2-mediated H3K27me3 levels and genomic distribution provide unique insights as to how transcriptionally active states are converted to repressive chromatin regions. CTR9 silencing results in the loss of imprinted genes during preimplantation development in mice (Zhang et al., 2013), which may cause genome instability. Loss-of-function mutations of CTR9 were recently discovered in Wilms tumors and mutations resulted in an in-frame deletion of exon 9. Whether H3K27me3 is elevated in CTR9-mutated Wilms tumors, and whether CTR9-mutant expressing tumors are sensitive to EZH2 inhibitors, awaits investigation. The new function of CTR9 in regulating PRC2-repressive H3K27me3 domains opens new avenues for understanding the biological functions of CTR9 in development and epigenetic therapies targeting the PRC2 complex in human cancers.

## MATERIAL & METHOD

### Cell lines and Cell Culture

HEK293T, MCF7, T47D, HS578T, MDA-MB-231 and MDA-MB-468 cell lines were obtained from American Type Culture Collection (ATCC) and maintained in Dulbecco’s modified medium (DMEM) (Gibco) supplemented with 10% fetal bovine serum (FBS) (VWR) and 1% Penicillin-Streptomycin (P/S) (Gibco). BT474, ZR-75 cell lines were obtained from ATCC and maintained in RPMI 1640 medium (Gibco) supplemented with 10% FBS and 1% P/S. MCF7-tet-on-parental cells and MCF7-tet-on-shCtr9 cells were generated previously (Zeng and Xu, 2015) and maintained in DMEM supplemented with 10% FBS and 1% P/S. All cells were cultured at 37℃ in a humidified atmosphere containing 5% CO_2_.

### 3D spheroids formation

5,000 – 30,000 cells of MCF7-shControl/shCtr9#3/shCtr9#5 were fully resuspended in 200 – 250 µl of DMEM media and seeded in 96-well round bottom plates with ultralow attachment. (Corning, Product# 4515) 3D spheroid cultures were grown at 37 °C up to 4 d in a humidified atmosphere with 5% CO_2_.

### Cloning and Plasmids

cDNAs for human Ctr9 (NM_014633.5), JARID2 (NM_004973.4) and KDM6A (NM_001291415.1) in pCMV-SPORT6 vector were purchased from Open Biosystems. The cDNAs were amplified using primers containing N-terminus Flag-tag and subcloned into pHAGE-PGK-GFP-IRES-LUC-W (Addgene plasmid # 46793) vector by replacing the luciferase-ORF with cDNAs encoding Ctr9, or JARID2 or KDM6A by applying the In-Fusion HD Cloning Plus Kit (Invitrogen) according to the manufacturer’s protocol.

### Virus packaging, infection and preparation of stable knockdown cell line

Stable knockdown or overexpression cell lines were generated by lentivirus infection. The virus packaging vectors pME-VSVG and psPAX2 were purchased from Open Biosystems. For lentivirus packaging, 4 μg pME-VSVG, 4 μg psPAX2 or 8μg lentiviral shRNA expression vectors or protein overexpression vectors were co-transfected into HEK293T cells cultured in one 10-cm dish using transIT-LT1 reagent (Mirus Bio) according to the manufacturer’s protocol. Medium was replaced with fresh DMEM supplemented with 10% FBS/ 1%P/S 12 hours after transfection. The supernatant containing the virus particles was collected by centrifugation (1500 rpm, 5 minutes) and subsequent filtered through a 0.45 μm syringe filter (Thermo Scientific) 48 hours after transfection. Approximately 1/5 volume of Lenti-X concentrator (Clonetech) was added to concentrate the virus tilter overnight at 4℃. For infection, 1 mL of virus was mixed with 1 mL of fresh cell culture medium, and polybrene was added at a final concentration of 8 μg/mL in order to increase the infection efficiency. After overnight infection, the culture medium was changed. Cells were infected overnight, followed by changing of culture medium. Cells were selected with 2 μg/mL puromycin for at least one week to generate stable cell lines.

### Preparation of the chromatin fraction

Cells were harvested after trypsinization. After washing with 1X PBS, two to five volumes of lysis buffer were added to the cell pellet [10mM HEPES pH7.4, 10mM KCl and 0.05% NP-40, 1X protease inhibitor cocktail (Sigma-Aldrich), phosphatase inhibitor (1mM NaVO4) and deacetylase (5mM TSA, Sigma-Aldrich)], and then incubated on ice for 20 mins. Nuclei pellets were separated by centrifugation at 14,000 rpm at 4℃ for 10 mins. Subsequently, nuclei pellets were washed once with lysis buffer, resuspended in two to five volumes of low salt buffer [10mM Tris-HCl pH7.4, 0.2mM MgCl_2_, 1X protease inhibitor cocktail phosphatase inhibitor (1mM NaVO4) and deacetylase (5mM TSA), and 1% Triton X-100], and incubated on ice for 15 minutes. Chromatin fractions were separated by centrifugation at 14,000 rpm at 4℃ for 10 mins, resuspended with two to five volumes of 0.2N HCl, and incubated on ice for 20 minutes. After centrifugation, the supernatant containing chromatin associated proteins was neutralized with an equal volume of 1M Tris-HCl pH 8.0.

#### Histone extraction and purification

Cells were harvested after trypsinization. After washing with 1X PBS, cell pellets were resuspended in two volumes of lysis buffer [50mM Tris-HCl pH7.4, 150mM NaCl, 10% Glycerol and 0.05% NP-40]. Protease Inhibitors (1X protease inhibitor cocktail) and HDAC inhibitor (10mM sodium butyrate) were added before use. The cell pellets were incubated on ice for 30 mins, followed by a brief sonication. After 15 mins of centrifugation at 13,000 rpm at 4°C, the supernatant was saved as whole cell lysate and the pellet was used for histone extraction. Pellets were washed twice using NIB buffer [10mM Tris-HCl pH7.5, 2mM MgCl_2_, 3mM CaCl2 and 1% NP40] containing 100mM NaCl and once with NIB buffer containing 400mM NaCl. The pellet was finally resuspended in NIB buffer (400mM NaCl) without NP40. For acid extraction of histones, double volume of 0.2N HCl was added and incubated overnight at 4℃. After centrifugation at 13,000 rpm for 15 mins at 4℃, solubilized histone in supernatant was dialyzed in ddH_2_O overnight at 4℃ using 10K MWCO Dialysis Tubing (Thermo). Histones were lyophilized and dissolved in ddH_2_O.

#### Liquid Chromatography and Quantitative Histone Mass Spectrometry (LC-MS/MS)

##### Chemical derivatization of histones and tryptic digestion

Take 25 μg of purified histone samples and dissolve them in 100μL 100mM TEAB buffer (pH 8.0). Add 4μL 4% ^13^CD_2_O (w/v) and 4μL 600 mM NaBD_3_CN to the samples and vortex them at room temperature for 1h to label the free and mono-methylated lysine with heavy isotopic methyl. Terminate the reaction by acidifying the sample with TFA. Transfer the samples to 10K MWCO ultracentrifuge tube (Millipore) and centrifuge the samples for 15 min at 14,000 g at 4℃ to remove the reaction reagents. Continually add 200μL 100mM TEAB buffer in the ultracentrifuge tube and centrifuge the samples for 15 min at 14,000 g at 4℃ to wash histone sample for 3 times. Add 100μL 100mM TEAB buffer and 1μg trypsin in the ultracentrifuge tube and put the samples at 37℃ incubators for 16 h. Collect the digested sample by centrifuging the samples for 15 min at 14,000 g at 4℃. Wash the ultracentrifuge tube two times with 100μL 100mM TEAB buffer and combine the follow-through with digested sample and dry down. Resolve the dry down sample with 100μL 50mM TEAB buffer and add 15μL 25% propionic anhydride buffer (v/v in ACN) in the sample, add 10-15 μL 28% NH_4_OH to keep the pH of the sample reaction to be around 8.0. Put the sample tube on vortex for 20 min to label the N-terminal of digested histone peptides with propionyl. After propionylation, the sample was desalted with Sep-Pak cartridge (Waters) and lyophilized.

##### LC-MS/MS for histone modification

Lyophilized histone peptides were resuspended in 0.1% formic acid (FA) and analyzed on a Dionex U3000 ultra performance liquid chromatography system coupled to a Q-Exactive HF quadrupole orbitrap mass spectrometer (Thermo Fisher Scientific). Peptide sample was injected (2 μl). A Waters BEH 300Å C18 reversed phase capillary column (150 mm x 75 µm, 1.7 µm) was used for separation. Water with 0.1% FA and acetonitrile with 0.1% FA were used as mobile phases A and B, respectively. The flow rate was set to 0.300 µL/min. 2 µL of sample was injected onto the column and separated over a 120-minute gradient as follows: 0-1 min 3-10% B; 1-90 min 10-35% B; 90-92 min 35-95% B; 92-102 min 95% B; 102-105 min 95-3% B; 105-120 min 3% B. The data was acquired under data dependent acquisition mode (DDA, top 20). Mass spectrometric conditions were as follows: spray voltage of 2.8 kV, no sheath and auxiliary gas flow; heated capillary temperature of 275°C, normalized high-energy collision dissociation (HCD) collision energy of 33%, resolution of 120, 000 for full scan, resolution of 60, 000 for MS/MS scan, automatic gain control of 2e5, maximum ion injection time of 100 ms, isolation window of 1.6, and fixed first mass of 110 m/z

##### Data analysis and relative quantification of histone PTMs

Due to histones possess multiple post-translational modifications and after heavy isotopic dimethylation and propionylation they are even highly modified and may contain several isoforms for same peptide sequence with different modifications. It is challenging to identify all different forms of histone modification peptides. Here in this work, the most frequently observed modifications that is the mono-methylation, di-methylation, tri-methylation, and acetylation on lysine residue of histone H3 were analyzed. The same histone peptide possess different modifications can firstly be identified according to their difference in mass over charge (m/z). Then the peptide can be additionally distinguished according to their retention time difference on the RP-HPLC column (trimethylated peptide ≈ unmodified (possesses two heavy isotopic methyl), mono (possesses one heavy isotopic methyl), and di-methylated peptide < acetylated peptides. To relatively quantify the abundance of histone PTMs, we used the area of each identified peptide peak in the MS chromatogram for comparison. Normally same peptide may have different charge state ions in LC-MS analysis and we only choose the highest intensity ions to measure their peak area. The total peak area of a histone peptide with all different PTM forms is regarded as 100%, and the percentage of each PTM the peptide is calculated by dividing the area of the PTM peak area by the total peak area. To distinguish histone peptide isoform, we also investigate the MS/MS spectrum to calculate the ratios of b and/or y ions that were different between the two or more peptide isoforms and the ratio is used to continually calculate the relative quantity of the peptide isoforms.

### Preparation of nuclear lysates

Nuclear lysates were prepared as described (Michael, H. K., et.al., 2010) and used for co-immunoprecipitation and peptide pulldown experiments. Cells were harvested by scraping in ice-cold PBS and extracted at 4℃ in buffer containing 50mM Tris-HCl pH7.5, 5mM EDTA, 250mM NaCl and 0.1% NP-40 supplemented with protease and phosphatase inhibitors for 30 mins. After centrifugation at 13,000 rpm at 4℃ for 1 hour, the supernatant was collected and mixed with two volumes of buffer containing 50mM Tris-HCl pH7.5, 5mM EDTA, 100 mM NaCl, 0.1% NP-40 and 10% glycerol.

### Co-Immunoprecipitation using nuclear extract

Co-IP was performed as previously described (Malte, B., et.al., 2016). Briefly, immunoprecipitations were performed in IP buffer [50mM Tris-HCl pH7.5,150mM NaCl, 2mM MgCl_2_, 0.5% NP-40 and 10% Glycerol] supplemented with protease and phosphatase inhibitors before use. Approximately 1.5 - 2 mg nuclear protein extract, as quantified by a Bradford assay, was mixed with 5 μg of antibody and 50 μl of protein A magnetic Dynabeads (Invitrogen, washed previously 3X in IP buffer) per IP reaction in 750 μl total volume. Beads were washed three times with IP buffer, and once with PBST the following day. Proteins were eluted in 75 μl 2X SDS loading buffer with 50nM DTT and heated at 95℃ for 15 mins before loading on an SDS-PAGE gel.

### Peptide pulldown using nuclear extract

Peptide pulldown was adapted from protocol described in (Malte, B., et.al., 2016). Lysine methylated peptide with a C-terminal biotin tag ATKAAR-Kme3-SAPSTGGVKKPHRYRPG-GGK(Biotin)-NH_2_, and the corresponding non-modified peptide were synthesized by Active Motif^®^. For each peptide pull down, 50 μg of magnetic streptavidin beads (Medchem Express) were incubated with 5 μg of peptide in 500 μl of binding buffer [50mM Tris-HCl pH7.5, 150mM NaCl, 1mM EDTA and 1% NP40] for three hours, rotating at room temperature. Peptide-bound beads were washed three times in wash buffer I [50mM Tris-HCl pH7.5, 150mM NaCl, 1mM EDTA and 0.05% Triton X-100], twice in wash buffer II [50mM Tris-HCl pH7.5, 150mM NaCl and 1mM EDTA]. Samples were vortexed twice with 50 μl U/T buffer (6M urea, 2M thiourea, 150mM NaCl, 30mM biotin in 10mM HEPES pH 8.0) at room temperature for 10 min and at 95℃ for 15 min. The two eluates were combined for mass spectrometry analysis

### Western blotting

Cells were harvested after trypsinization, washed with Dulbecco’s phosphate buffer saline (DPBS) (Life Technologies), and lysed in lysis buffer [50 mM Tris-HCl pH 8.0, 400 mM NaCl, 10% glycerol, 0.5% Triton X-100 and 1 x protease inhibitor cocktail (Sigma-Aldrich)]. After a brief sonication, total lysate was centrifuged, and the supernatant was quantified using the BioRad Protein Assay (BioRad). Approximately 30 μg protein was resolved by SDS-PAGE. Proteins were transferred to a nitrocellulose membrane for 1.5 hours at 350 mA. Membranes were blocked with 5% nonfat milk or 5% BSA at room temperature for 1 hour and incubated overnight with diluted primary antibody at 4 °C. Membranes were then washed and incubated with HRP-conjugated goat-anti-rabbit or mouse IgG secondary antibody for 1 hour at room temperature. Membrane was incubated with enhanced chemiluminescence reagents (Thermo Scientific) followed by exposure to X-ray films.

### ELISA for Histone modifications

ELISA measurement of specific Histone H3 modification was performed in using Histone H3 Modification Multiplex Assay Kit (abcam) according to manufacturer’s protocol. In brief, extracted histone mixture in Antibody Buffer (provided in kit) was aligned to wells coated with specific Histone H3 modification antibodies. After incubation at 37℃ for 2hrs, solution in wells was removed and wells were washed 3X in wash buffer (provided in kit). Diluted detection antibody solution (provide in kit) were added and incubate at room temperature for 1hr. After 2x brief wash in wash buffer, developer solution was added and incubate at room temperature for 2-10 mins until wells sufficiently turn into blue while blank wells with no histone extract added remain transparent. Stop the reaction by adding the stop solution (provided in kit) and the wells will turn into yellow. The absorbance of each well was read on a microplate reader at 450 nm with an optional reference wavelength of 655 nm. Absorbance output among different samples will first eliminate the background noise measured by the blank wells and subsequently normalized by corresponding absorbance in control wells coated with H3-total antibody.

### Cell proliferation and two-dimensional (2D) colony formation assays

For cell counting-based proliferation assays, 1 × 10^5^ cells were seeded into six 3.5 cm petri dishes for compound treatment. MCF7-tet-on–shCtr9 cells were pretreated with vehicle or 500 ng/mL Dox for 5 days before seeding to 3.5 cm petri dishes. Media were changed every 48 hours. Cells were trypsinized and counted after Trypan blue exclusion using an automated cell counter (Bio-Rad).

For 3-(4,5-dimethylthiazol-2-yl)-2,5-diphenyltetrazolium (MTT)-based (Sigma-Aldrich) proliferation assays, 2 × 10^3^ cells were seeded into 96-well plates for compound treatment. 15 μl MTT (5 mg/mL in DPBS) was added to the cells followed by incubation at 37°C for 1 h. After removing cell culture medium, 50 μL of DMSO was added. The absorbance of the color substrate was measured with a 540-nm filter on a VictorX5 microplate reader (Perkin Elmer), and data were plotted and analyzed using GraphPad Prism 7 software (GraphPad Software, Inc.).

For 2D colony formation assays, 1000 cells were seeded into 3.5 cm petri dishes for compound treatment, and media was refreshed every 4 days. After treatment with compounds for 12-15 days, cells were washed with DPBS, fixed with 4% formaldehyde for 10 min at room temperature, and stained with 0.05% crystal violet for 30 min at room temperature. Images were taken on a Leica inverted microscope using the Leica Application Suite. Colony numbers were counted using ImagePro software.

### Senescence-associated β-galactosidase staining

Cells were fixed in a 2% formaldehyde / 0.2% glutaraldehyde solution for 5 min, and then stained with an X-Gal staining buffer (20mM Citric Acid pH 6.0, 40mM Dibasic-Na_3_(PO_4_)_2_, 5mM K_4_[Fe(CN)_6_], 5mM K_3_[Fe(CN)_6_], 150mM NaCl, 2mM MgCl_2_ and 1mg/ml X-gal in DMF) overnight at 37°C. After washing with PBS twice, cells were imaged on a Leica inverted microscope using the Leica Application Suite.

### Cell cycle analyses using flow cytometry

For PI staining-based cell cycle analysis, cells were harvested after trypsinization, fixed in cold 95% ethanol, and washed in PBS. The fixed cells were then resuspended in propidium iodide staining solution (200 μg/mL RNase A, 50 μg/mL propidium iodide, 0.1% [v/v] Triton X-100 in PBS +1% BSA) and incubated overnight at 4°C. Flow cytometry analyses were performed at the University of Wisconsin Flow Cytometry Laboratory and data were analyzed using FlowJo software (FlowJo, LCC).

Edu staining-based cell proliferation analysis was performed by using the Click-iT^TM^ Plus Edu Flow Cytometry Assay Kit (Invitrogen) according to manufacturer’s protocol. In brief, cells were labeled with 10 μM Edu staining buffer in culture medium for 2 hours. After a brief wash with 1% BSA in DPBS, cells were harvested by centrifugation. Cells were then fixed in fixative buffer (provided in kit) at room temperature for 15 mins. After washing with 1% BSA in DPBS twice, cells were resuspended in saponin-based permeabilization buffer (provided in kit) at room temperature for 15 mins. 500 μl Click-iT^TM^ Plus reaction cocktail was added to each sample, followed by incubation of the reaction mixture in dark at room temperature for 30 mins. Cells were washed with permeabilization buffer, wash buffer (provided in kit), and then subjected to flow cytometry analysis.

### Annexin V and PI Staining by Flow Cytometry

Collect 1-5 × 10^5^ trypsinizied cells by centrifugation. Wash cells 1X with cold PBS and carefully remove the supernatant. Re-suspend the cells in Binding buffer [10 mM Hepes pH 7.4, 140 mM NaCl and 2.5 mM CaCl2] at a concentration of ∼1 × 10^6^ cells/mL. After a brief centrifugation, cells were resuspended and incubated for 10 min with 0.5 μg/mL Annexin V-FITC and 2 μg/mL PI in 400 μL binding buffer. The cells were immediately placed on ice and analyzed by flow cytometry. Cell fragments were removed by morphological gating. Cells negative for Annexin V-FITC and PI were considered viable, Annexin V-FITC positive and PI negative considered apoptotic, and Annexin V-FITC positive and PI positive considered necrotic.

### Cytotoxicity assay of 3D spheroids

Assay was performed according to manufacturer’s instruction (Invitrogen, Cat# L3224). In brief, optimized concentration of Calcein AM and Ethidium homodimer-1 as well as 1 µM Hoechest 33342 were added in DMEM media and incubated with spheroids at 37℃ for 30 mins. After incubation, those 3D spheroids were immediately subjected to confocal imaging (Nikon W1 confocal) in a live cell incubating chamber. The area of 3D spheroids was measured using NIS-A1R Advanced Research Imaging Software (Nikon - Mager Science) with nuclei annotation.

### Immunofluorescence staining of H3K27me3

MCF7-tet-on-shCtr9 cells were seeded on glass bottom 3.5cm petri dishes and cultured in DMEM supplemented with 10% FBS in the absence or presence of 500 ng/mL Dox. Cells were fixed in 4% formaldehyde for 15 mins, and then washed with DPBS three times. Subsequently, cells were permeabilized in 0.3% Triton X-100 in PBS for 10 mins, blocked with 3% BSA in PBST (PBS + 0.1% Triton X-100) for 1hr, and incubated with H3K27me3 primary antibody (Cell signaling technology, #9733:C6B11) at room temperature for 2 hours. Cells were then washed with PBST, followed by incubation with secondary antibody (Cy5-goat anti-rabbit, 1:250; Bethyl) for 30 mins at room temperature. After being washed twice in PBST, cells were incubated with 50 nM Alexa Fluor 555 Phalloidin (Cell Signaling Technology) and 1 μg/ml of Hoechest 33342 (Cell Signaling Technology) at 37℃ for 15 mins and washed twice in DPBS. Fluorescence was detected using a Nikon A1R confocal microscope at appropriate wavelengths at the UW imaging core. And signal intensity was analyzed in NIS-A1R Advanced Research Imaging Software (Nikon - Mager Science)

### Quantification of H3K27me3 intensity using flow cytometry

The Dox treatment of MCF7-tet-on-shCtr9 cells was performed as previously described above. Cells were harvested after trypsinization, washed in PBS, and fixed in 4% paraformaldehyde for 15 mins. Subsequently, cells were permeabilized with 0.3% Triton X-100 in PBS for 10 mins, blocked with 3% BSA in PBST (PBS + 0.1% Triton X-100) for 1 hour, and incubated with Cy5 conjugated H3K27me3 antibody (abcam) in 1% BSA in PBST for 1hr. After washing with PBST, cells were subjected to flow cytometry analysis.

### Total RNA and Ribosome associated RNA preparation and real-time quantitative PCR (RT-qPCR)

Total RNA was extracted using the HP Total RNA Kit (VWR Scientific) according to the manufacturer’s protocol. Ribosome associated RNA in cycloheximide pretreated cells were similarly prepared as described in (Myriam, H., et.al., 2014) by using 40S ribosomal protein S3 antibody (Cell signaling technology) to perform immunoprecipitation. Dynabeads^TM^ Protein A/G (Invitrogen) was used to enrich ribosome-RNA complex at the next day. Eluted RNA was further cleaned up by using miRNeasy Micro Kit (Qiagen). 2μg of total RNA or ribosome associated RNA were reverse transcribed using RevertAid First Strand cDNA Synthesis kit (Thermo Fisher Scientific). Quantitative PCR was performed using Fast Start Universal SYBR Green Master Mix (Roche) on a BioRad CFX-96 instrument. Primer sequences are summarized in Supplemental Table S3. Data were analyzed using the ΔΔCq method calculated by the CFX Manager Software (BioRad).

### Chromatin immunoprecipitation (ChIP)

Cells in 15-cm dishes were washed once with PBS before cross-linking with PBS containing 1% formaldehyde for 15mins at room temperature. Crosslinking was quenched with 0.125 M glycine for 5 minutes at room temperature before two washes with ice-cold PBS. Cells were scraped, harvested by centrifugation, and subjected to ChIP assays. Crosslinked cells were lysed with lysis buffer 1 (10 mM HEPES pH 7.0, 10 mM EDTA, 0.5 mM EGTA, 0.25% Triton X-100, supplemented with 0.5 mM PMSF before use) with rotation at 4 °C for 10 minutes. The crude nuclear pellets were collected by centrifugation at 1500 rpm for 4 minutes at 4 °C. The supernatant was discarded, and the chromatin was washed with lysis buffer 2 (10 mM HEPES pH 7.0, 200 mM NaCl, 1 mM EDTA, 0.5 mM EGTA, supplemented with 0.5 mM PMSF before use) for 10 minutes at 4 °C with rotation. Nuclear pellets were collected by centrifugation (1500 rpm, 4 °C, 4 minutes), resuspended in nuclear lysis buffer [50 mM Tris-HCl pH 8.1, 10 mM EDTA, 1% SDS, supplemented with 1 mM PMSF and 1 x protease inhibitor cocktail (Sigma-Aldrich) before use, and incubated on ice for 10 minutes. Chromatin was sheared to approximately 100bp-1000bp fragments by sonication in ice-water bath at 4 °C using a Branson Sonifier 450 with a microtip (40% amplitude, 3 seconds on, 10 seconds off, 3 minutes total pulse time). Sonicated chromatin was centrifuged at 15,000 rpm for 15 minutes at 10 °C, and concentration of nuclear proteins was determined using the BioRad Protein Assay (BioRad). Equal amounts of total nuclear proteins were used for ChIP. Nuclear proteins were supplemented with nuclear lysis buffer to achieve same final volumes between different samples, and then diluted 1:10 with dilution buffer (20 mM Tris-HCl pH 8.1, 150 mM NaCl, 2 mM EDTA, 1% Triton X-100, supplemented with 1 x protease inhibitor cocktail before use). Five percent of the chromatin fraction was removed and saved as input, and the rest was pre-cleared with a normal IgG control before incubating with the antibody of interest overnight at 4 °C.

On the following day, the immune complexes were incubated with Dynabeads^TM^ Protein A/G or Dynabeads^TM^ M-280 Sheep anti-Mouse IgG (Life Technologies) (beads were pre-washed with ChIP dilution buffer three times before use) while rotating at 4 °C for 2 hours. The immunoprecipitated materials were subsequently washed once with low salt wash buffer (20 mM Tris-HCl pH 8.1, 150 mM NaCl, 2 mM EDTA, 0.1% SDS, 1% Triton X-100), once with high salt wash buffer (20 mM Tris-HCl pH 8.1, 500 mM NaCl, 2 mM EDTA, 0.1% SDS, 1% Triton X-100), once with LiCl wash buffer (10 mM Tris-HCl pH 8.1, 0.25 M LiCl, 1 mM EDTA, 1% NP-40, 1% deoxycholate), and twice with TE buffer (10mM Tris-HCl pH 8.0, 1 mM EDTA pH 8.0). Each wash was done with rotation at 4 °C for 5 minutes followed by separation on magnetic stand. The immunoprecipitated material was eluted twice in freshly prepared elution buffer (1% SDS, 0.1 M NaHCO3) with shaking on a vortexer for 20 minutes at room temperature. The eluted and input materials were then digested with proteinase K (200 μg/ml final concentration) at 55 °C for 2 hours. The crosslinking materials were reversed by incubating at 65 °C in a hybridization oven overnight. DNA was purified using a Qiagen PCR Purification Kit. The ChIPed DNA was then analyzed by qPCR using the SYBR Green detection chemistry and the location-specific primers in Supplemental Table II. Fold change was determined by normalizing the ChIPed DNA signal to the input DNA signal.

### ChIP-seq library preparation

Prior to ChIP-seq library preparation, the concentration and size distribution of the ChIPed DNA samples was determined using a Qubit Fluorometer (Thermo Fisher Scientific) and the Agilent High Sensitivity DNA Kit (Agilent Technologies), respectively at the Sequencing Facility at the University of Wisconsin-Madison Biotechnology Center. Approximately 10-25 ng of ChIPed DNA from each condition was used to generate the ChIP-seq library using the Ovation Ultralow System V2 1-16 Kit (NuGEN Technologies), according to the manufacturer’s protocol. Briefly, the DNA was end-repaired and ligated to Illumina sequencing adaptors. The ligated DNA was purified using Agencourt RNAClean XP beads (Beckman Coulter). A subsequent PCR amplification step (15 cycles) was performed to add linker sequence to the purified fragments for annealing to the Genome Analyzer flow-cell. Following PCR amplification, the library was separated on a 2% agarose gel (120 V, 1.5 hours) to select a narrow range of fragment sizes, and bands between 200-500 bp were excised. The library was purified from the excised agarose gel using the Qiagen MiniElute PCR Purification Kit following the manufacturer’s protocol.

Quality control for the size, purity, and concentration of the final ChIP-seq libraries was performed at the Sequencing Facility at University of Wisconsin**-**Madison Biotechnology Center. Qualified libraries were deep sequenced using an Illumina HiSeq 2000 per the manufacturer’s instructions at the University of Wisconsin Biotechnology Center.

### ChIP-seq data analysis

ChIP-seq reads were aligned to human genome (hg38) by Bowtie (version 1.1.2) with command line options ‘--quiet -q -v 2 -a --best --strata -m 1 --phred33-quals -S’. We removed reads that were unmapped or PCR/optical duplicates and those that did not pass platform/vendor quality controls. ChIP-seq peaks were called by MACS (version 2.1.0.20151222) with a q-value cutoff of 0.05. Narrow peaks were called for H3K4me3 and broad peaks were called for H3K27me3 and H3K36me3. Peaks that overlapped with ENCODE’s Blacklisted Regions (https://www.encodeproject.org/annotations/ENCSR636HFF/) were removed. An overlap was defined as having at least one base-pair in common. We applied the same definition when comparing peaks from different conditions and a unique peak in one condition required that it did not have any overlap with peaks from the other condition in comparison. ChIP-seq signals were calculated by MACS as per million reads for fragment pileup profiles in bedGraph format and converted to bigWig format.

To prepare heatmaps of H3K27me3 signals, a ‘max-center’ was picked within each H3K27me3 peak by selecting the genomic locus with the highest H3K27me3 signal. If there were multiple loci with the highest signals, their geometric center will be defined as the ‘max-center’. Each max-center’s upstream and downstream 2kb regions were divided into 200 bins and each bin had an equal width of 10 bp. Average H3K27me3 signals were calculated for each bin. Bin signals across all the peaks within each cluster were further averaged and shown above the heatmap for the corresponding cluster.

We used GENCODE v27 basic gene annotation on the reference chromosomes only (https://www.gencodegenes.org/human/release_27.html) to define genomic locations and types of genes and their transcripts. The whole human genome were divided into four types of regions: (i) ‘exon region’, representing exons of protein-coding transcripts; (ii) ‘intron region’, representing introns from protein-coding transcripts and do not overlap with any ‘exon regions’ (Note that a gene may have multiple alternatively spliced isoforms, i.e., transcripts, and one isoform’s exon may overlap with another isoform’s intron); (iii) ‘promoter regions’, defined as the 5kb upstream region of protein-coding transcript’s Transcription Start Site (TSS) and do not overlap with any ‘exon region’ or ‘intron region’; (iv) ‘intergenic region’, representing the remaining genomic regions that do not belong to any of the above three types of regions. We further divided ChIP-seq peaks by these four types of regions: (i) ‘promoter peak’: peaks overlapped with ‘promoter region’; (ii) ‘intron peak’: peaks that overlapped with ‘intron region’ and do not overlap with any ‘promoter region’; (iii) ‘exon peak’: peaks overlapped with ‘exon region’ and do not overlapped with any ‘promoter region’ or ‘intron region’; (iv) ‘intergenic peak’: peaks that does not belong any of the other three peak categories. Peaks from the first three categories were also considered as ‘genic peak’.

To study H3K27me3 signals around TSS, we selected protein-coding transcripts that gained H3K27me3 peaks in genomic region 5 kb around their TSS ([TSS - 5kb, TSS + 5kb]) after Ctr9 knockdown. For these selected transcripts, we divided their [TSS-5kb, TSS+5kb] region into 100 bins, where each bin has an equal width of 100 bp. H3K27me3 signals were calculated for each bin and averaged across all transcripts.

We used GENCODE v27 basic gene annotation on the reference chromosomes only to define genomic locations and types of genes and their transcripts. The whole human genome were divided into four types of regions: (i) ‘exon region’, representing exons of protein-coding transcripts; (ii) ‘intron region’, representing introns from protein-coding transcripts and do not overlap with any ‘exon regions’ (Note that a gene may have multiple alternatively spliced isoforms, i.e., transcripts, and one isoform’s exon may overlap with another isoform’s intron); (iii) ‘promoter regions’, defined as the 5kb upstream region of protein-coding transcript’s Transcription Start Site (TSS) and do not overlap with any ‘exon region’ or ‘intron region’; (iv) ‘intergenic region’, representing the remaining genomic regions that do not belong to any of the above three types of regions. We further divided ChIP-seq peaks by these four types of regions: (i) ‘promoter peak’: peaks overlapped with ‘promoter region’; (ii) ‘intron peak’: peaks that overlapped with ‘intron region’ and do not overlap with any ‘promoter region’; (iii) ‘exon peak’: peaks overlapped with ‘exon region’ and do not overlapped with any ‘promoter region’ or ‘intron region’; (iv) ‘intergenic peak’: peaks that does not belong any of the other three peak categories. Peaks from the first three categories were also considered as ‘genic peak’.

For Figure S4D, 240 Ctr9-regulated genes were identified in previous published work. [Zeng et al., 2016]. For two out of the 240 genes, *LOC100128881* and *LOC728342*, we could not map them to the current gene annotation and hence we analyzed the remaining 238 genes. Since a gene may contain multiple transcripts, we calculated individual transcript’s ChIP-seq signals and averaged them across all the transcripts from that gene. For H3K4me3, we calculated its signal over a transcript’s promoter, which was defined as the 5kb region upstream of a transcript’s TSS. For H3K27me3 and K3K36me3, we calculated their signals over a transcript’s promoter and a transcript’s genomic span (i.e., all exons and introns). UpSet plots were used to compare the number of genes with increased H3K27me3 signals and/or decreased H3K36me3 and H3K4me3 signals.

PRC2-EZH2 repressed genes were obtained from a recent study in mouse (*Foxc1* and genes listed in Figure 4a of [Hirukawa et al., 2018]. All of the 24 mouse genes can be mapped uniquely to human genes except that *Dbc1* was mapped to *BRINP1* and *CCAR2*, so we analyzed all of the 25 human genes. We defined gene body as the union of genomic spans (i.e., all exons and introns) of all the transcripts from that gene. We calculated H3K27me3 signals over the gene bodies of these 25 genes before and after Ctr9 knockdown.

### mRNA expression correlation analysis using published TCGA breast tumor RNA-seq data

For Figure S1F, complete correlation analysis and dot plotting were directly derived from the cBioPortal for Cancer Genomics (http://www.cbioportal.org/) by choosing RNA-seq datasets from [Ciriello et al., 2015]. 593 primary ER+ breast tumors with RNA-seq data were selected and Spearman correlation and corresponding p-values were recorded.

For Figure 5D, TCGA Breast Cancer clinical records and RNA-seq datasets were downloaded from Genomic Data Commons Data Portal (https://portal.gdc.cancer.gov).

Estrogen receptor (ER) status was determined by the entry ‘breast_carcinoma_estrogen_receptor_status’ in patient’s clinical record. There are 803 ER positive patients have primary tumor RNA-seq data. Upper Quantile normalized FPKM (FPKMUQ) were used to study gene expression levels. If a patient had multiple primary tumor RNA-seq datasets, we took the average of gene’s FPKMUQ across all the datasets. Pearson correlation coefficient was calculated between two genes’ log_10_(FPKMUQ). P-values were adjusted by Benjamini & Hochberg procedure to account for multiple hypothesis testing.

## DATA AVAILABILITY

The accession number for the ChIP-seq data reported in this paper is GEO: GSE133318. Source data have been provided separately with supplementary files.

## QUANTIFICATION AND STATISTICAL ANALYSIS

Statistical comparisons between two groups for RT-qPCR data and proliferation analyses were performed with Graphpad Prism software 7.0 using a paired two tails t-test. The sample size (n) is indicated in the figure legends and represents biological replicates. Details for sequence data analyses and statistical significance are described in the specific Method section.

## ACKNOWLEDGEMENTS

The authors thank Drs. Emery Bresnick, Peter Lewis and Lixin Rui for comments, Kristine Donahue for editing, and UWCCC flow cytometry center for technical support. The project is supported in part by NIH/NCI P30CA014520-UW Carbone Cancer Center Support Grant, and NIH R01 CA213293 and R01 CA236356 to W. X., and NIH S10RR029531, P41GM108538 grants to L.J. L.

## Contributions

W.X. and N.T.C. conceived the project, designed the experiments, analyzed the data and wrote the manuscript. N.T.C. performed the experiments with assistance from Y.D.W., J. H. performed the mass spectrometry experiments; P.L. and I.M.O. performed bioinformatics analyses; W.X. and L.J.L. directed and supervised all aspects of the study, all authors discussed the results and commented on the manuscript.

### Inventory of Supplemental Information

Key resources table

**Figure S1 (related to Figure 1).**
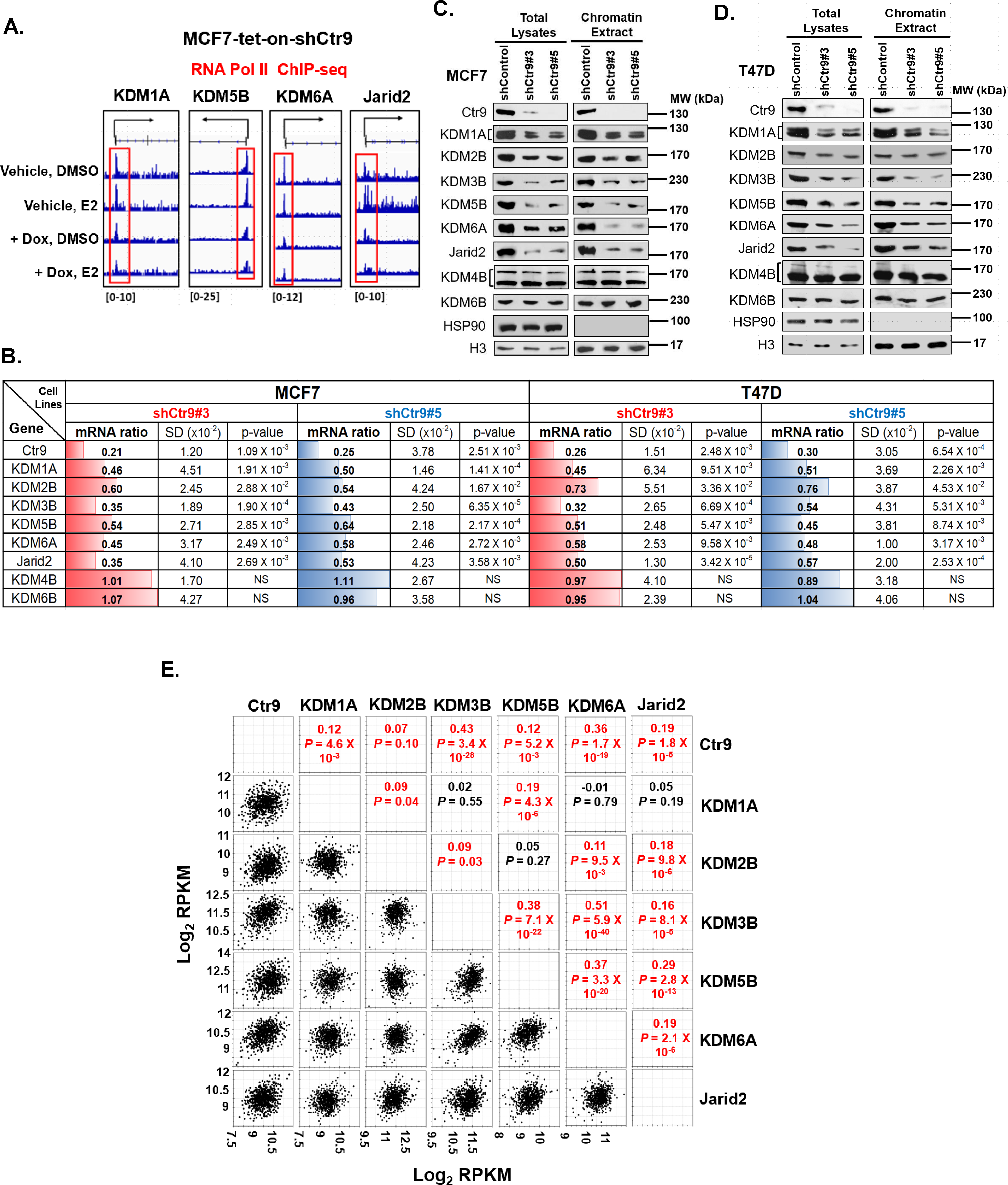
CTR9, but not other Paf1c subunits, regulate KDM protein expression in breast cancer cells. A. Representative genome browser views of RNA Pol II binding around the TSSs of KDM genes in MCF7-tet-on–shCtr9 cells treated with vehicle or Dox followed by DMSO or 10 nM E2 treatment. B. RT-qPCR analyses of mRNA levels of CTR9 and KDM genes in ER^+^ breast cancer cell lines MCF7/T47D expressing shControl, shCtr9#3 or shCtr9#5. Relative fold change in mRNA levels are represented as means ± SD (n = 3) and were normalized to the internal control β-Actin. P-values were calculated using a paired two tails t-test. C. Western blot of KDM proteins in total lysates or extracted chromatin fractions from MCF7-shControl, MCF7-shCtr9#3 or MCF7-shCtr9#5 cells. HSP90 and histone H3 served as loading controls for total lysates, and the chromatin extract, respectively. D. Western blot of KDM proteins in total lysates or extracted chromatin fractions from T47D-shControl, T47D-shCtr9#3 or T47D-shCtr9#5 cells. HSP90 and histone H3 served as loading controls for the total lysates, and the chromatin extract, respectively. E. Dot plot showing the correlation between CTR9 and KDMs at the mRNA level in 593 ER+ invasive breast tumors from TCGA (n = 593 biologically independent patient samples).Spearman correlation and corresponding p-values were shown.

**Figure S2 (related to Figure 2).**
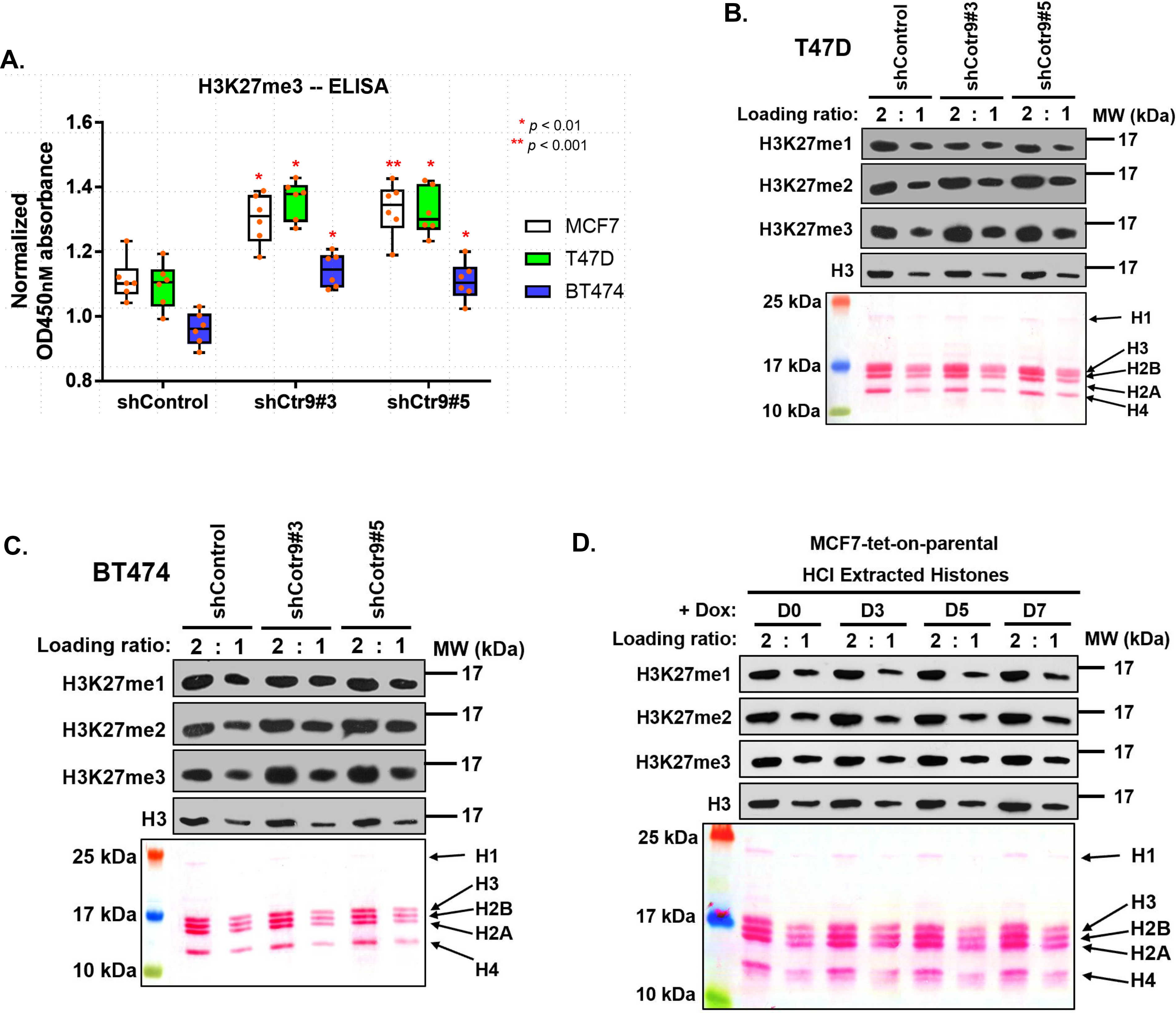
Depletion of CTR9, but not other Paf1c subunits, leads to the global increase of H3K27me3 in multiple breast cancer cell lines. A. ELISA analyses of H3K27me3 levels in MCF7, T47D and BT474 expressing shControl, shCtr9#3 or shCtr9#5. Data are represented as the mean ± SD (n = 6) and were normalized to respective total H3 levels. P-values were calculated using paired two tails t test. B. Western blot of H3K27me1/2/3 in T47D cells stably expressing shControl, shCtr9#3, or shCtr9#5 (top). Ponceau S staining of HCl extracted histones diluted two-fold for loading (bottom). C. Western blot of H3K27me1/2/3 in BT474 cells stably expressing shControl, shCtr9#3, or shCtr9#5 (top). Ponceau S staining of HCl extracted histones diluted two-fold for loading (bottom). D. Western Blot of H3K27me1/2/3 in MCF7-tet-on-parental cells in a time course Dox treatment (top). Ponceau S staining of HCl extracted histones in two-fold dilution for loading (bottom).

**Figure S3 (related to Figure 4).**
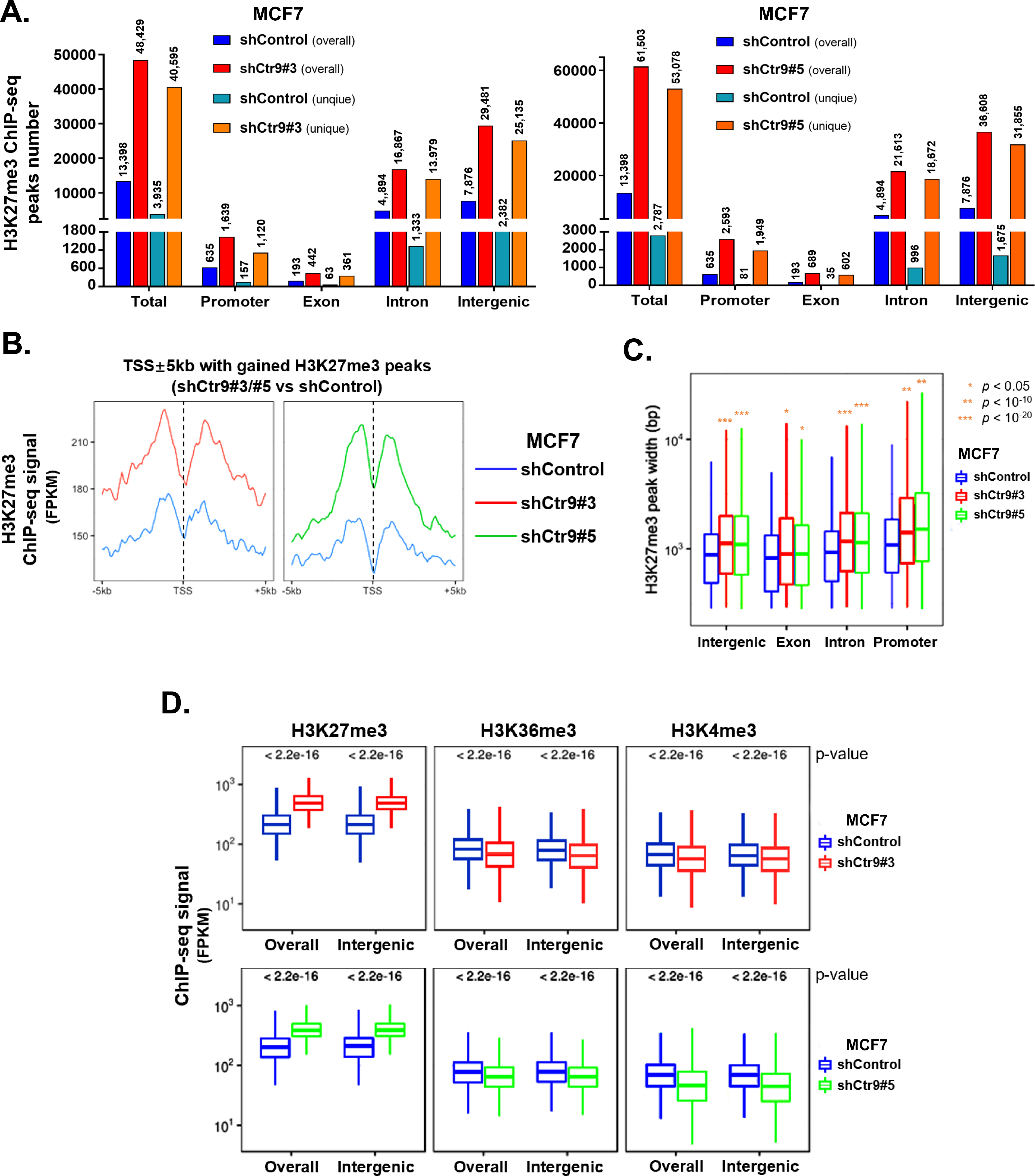
Stable depletion of CTR9 results in increased H3K27me3 peak numbers, intensity, and width. A. Summary of overall and unique H3K27me3 peak numbers in MCF7 cells stably expressing shControl, shCtr9s#3, or shCtr9#5. Peak numbers are classified as the total number of peaks, as well as the number of peaks in the promoter, exons, introns, or intergenic regions. B. Knocking down CTR9 using shCtr9#3 and shCtr9#5 induced a statistically significant increase of H3K27me3 and decrease of H3K4/K36me3 ChIP-seq signals across the genome and intergenic regions as compared to the shCtr9 control. P-values are shown on the top of each bar chart. C. Genome-wide increase of H3K27me3 ChIP-seq peak width when CTR9 was stably knocked down by shCtr9#3 and shCtr9#5. P-values were calculated using a one-sided Mann-Whitney U test. D. The average ChIP-seq profiles of H3K27me3 signal at those TSS ± 5kb regions with gained H3K27me3 peaks after CTR KD in MCF7 cells expressing shCtr9#3 (left) or shCtr9#5 (right) relative to MCF7 cells expressing control shRNA.

**Figure S4 (related to Figure 4).**
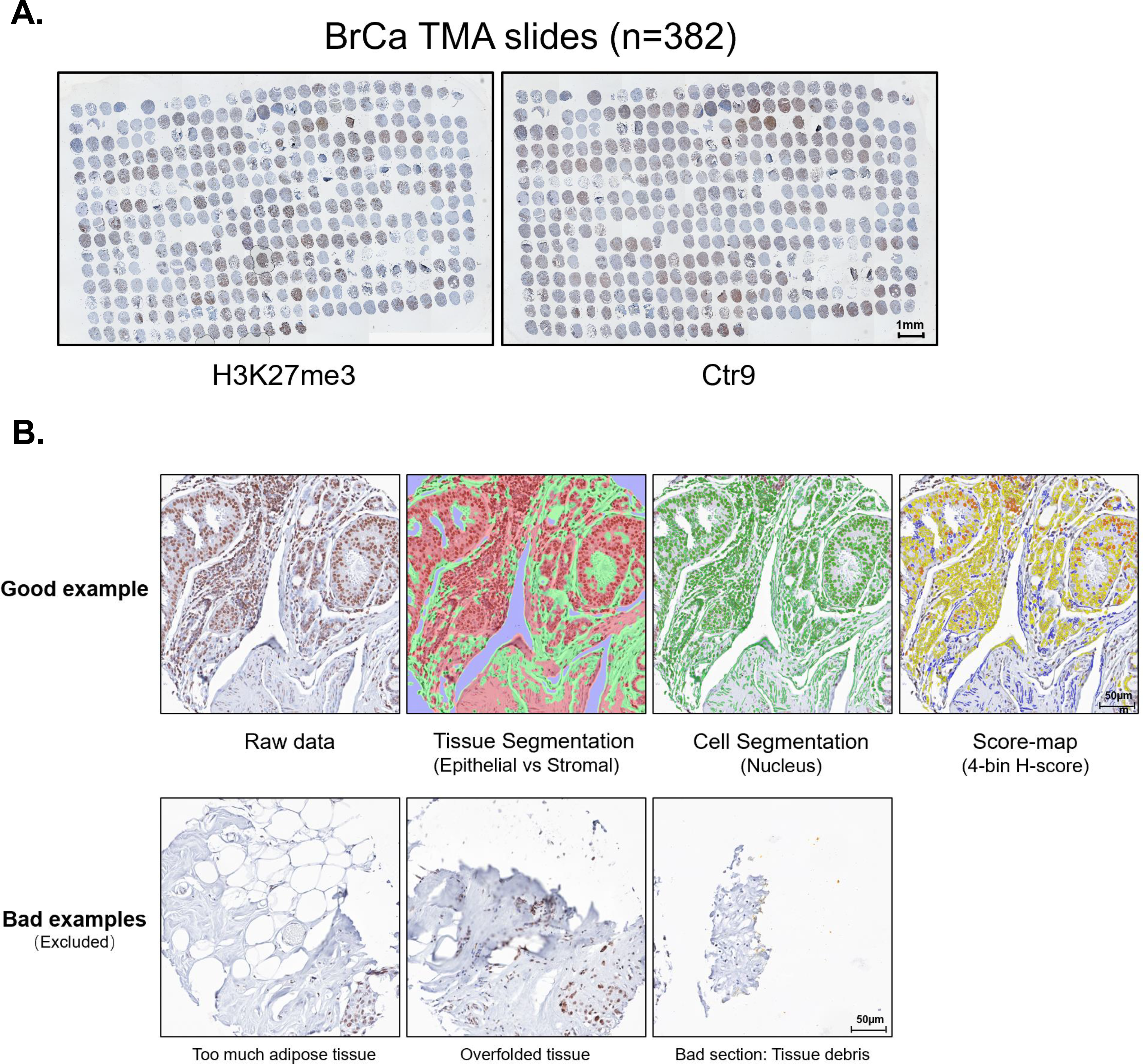
CTR9 has inverse correlation with H3K27me3 in breast cancer TMA samples. A. A general snapshot of breast cancer (BrCa) TMA slides (382 tumor cores) with H3K27me3 (left) and Ctr9 (right) IHC staining. B. Workflow for analyzing Ctr9 or H3K27me3 IHC staining in the nucleus of tumor epithelial cells and H-score grading in appropriate tumor cores. (Top) Representative bad samples of tumor cores within TMA slides that need to be excluded for correlation analysis. (Bottom)

**Figure S5 (related to Figure 1 and Figure 3).**
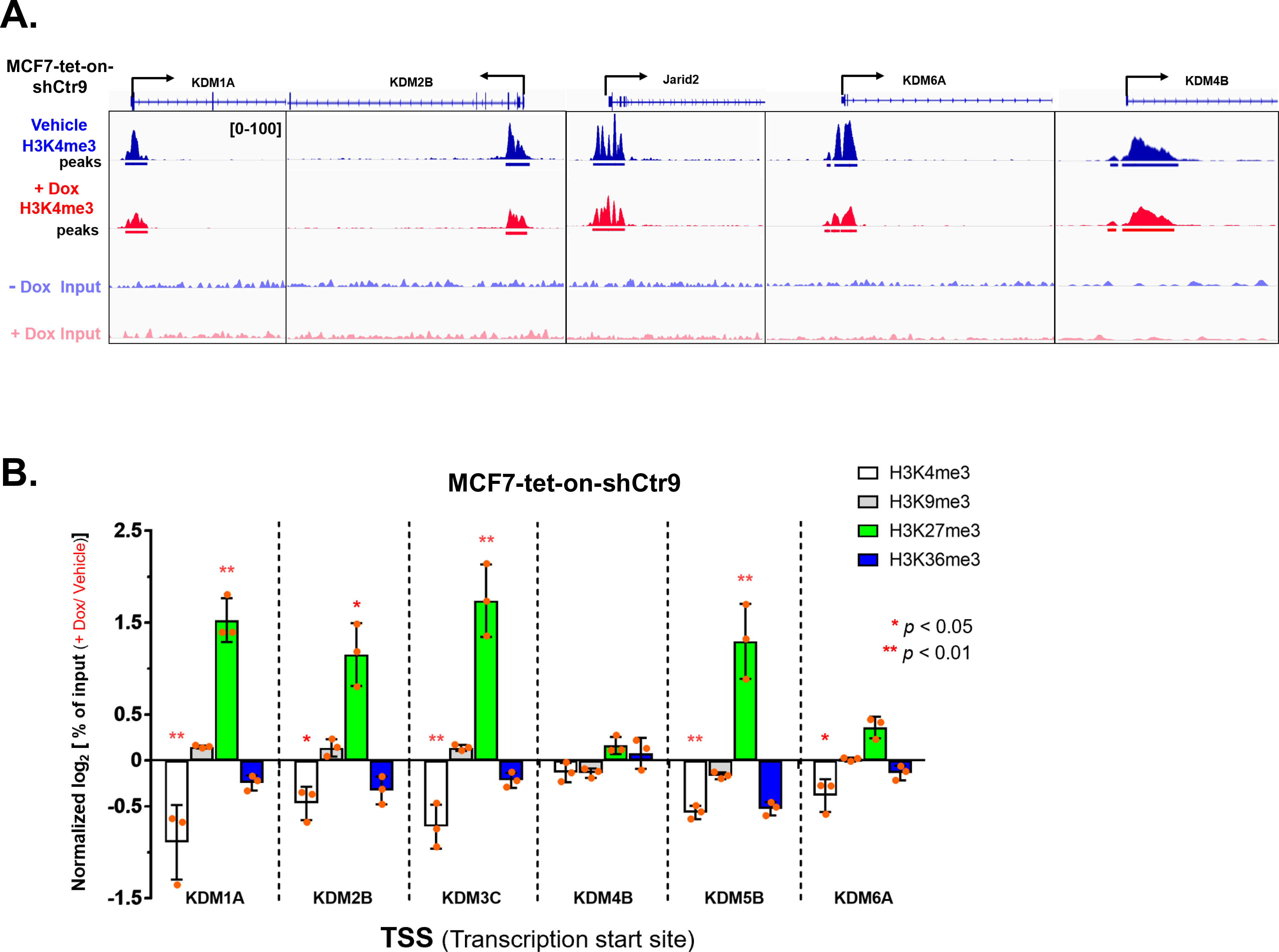
CTR9-regulated KDM genes are subjected to PRC2 and H3K27me3 regulation. A. Genome browser snapshots of H3K4me3 ChIP-seq profiles at the TSSs of CTR9-regulated KDM genes in MCF7-tet-on-shCtr9 cells treated with vehicle or Dox. KDM4B serves as a negative control. B. ChIP-qPCR analyses of H3K4me3, H3K9me3, H3K27me3, and H3K36me3 signals at the TSSs of KDM genes in MCF7-tet-on-shCtr9 cells treated with vehicle or Dox. The modified histone ChIP inputs were normalized to the total H3-ChIP input, and the ratios of the ChIP signal over input between Dox and vehicle treated group were log2 transformed (n=3). P-values were calculated using a paired two tails t-test.

**Figure S6 (related to Figure 5).**
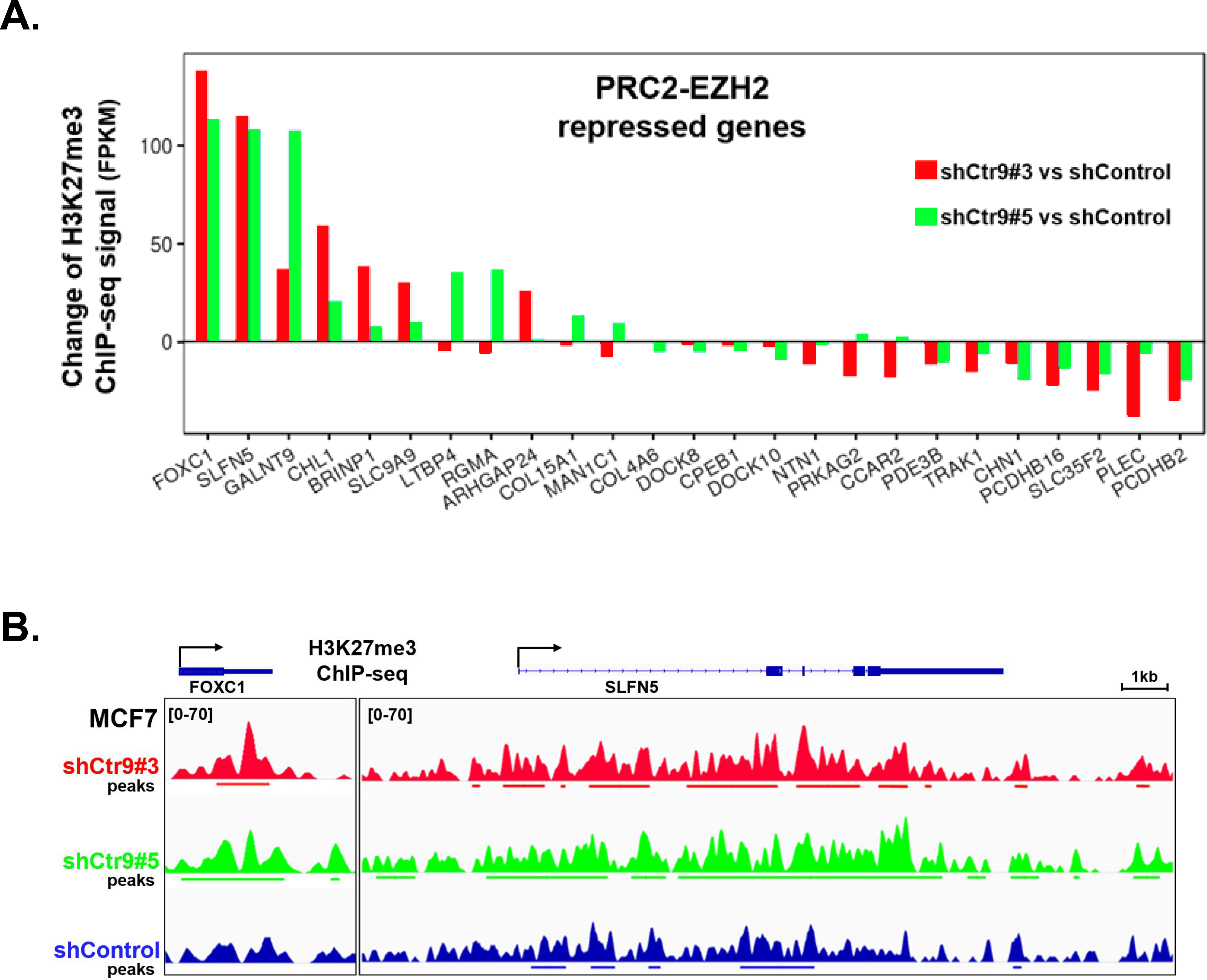
Expression of PRC2/EZH2 target genes are regulated by CTR9 in breast cancer cells. A. Change of H3K27me3 ChIP-seq signal on 25 human analogous genes after stable knockdown Ctr9 (shCtr9#3 or shCtr9#5 vs shControl). These 25 human homologous genes are derived from PRC2-EZH2 repressed genes in mouse. B. Representative genome-browser snapshots of H3K27me3 ChIP-seq profile at FOXC1 and SLFN5 genes with increased H3K27me3 ChIP-seq signal when Ctr9 was lost (shCtr9#3 or shCtr9#5 vs shControl).

**Figure S7 (related to Figure 6).**
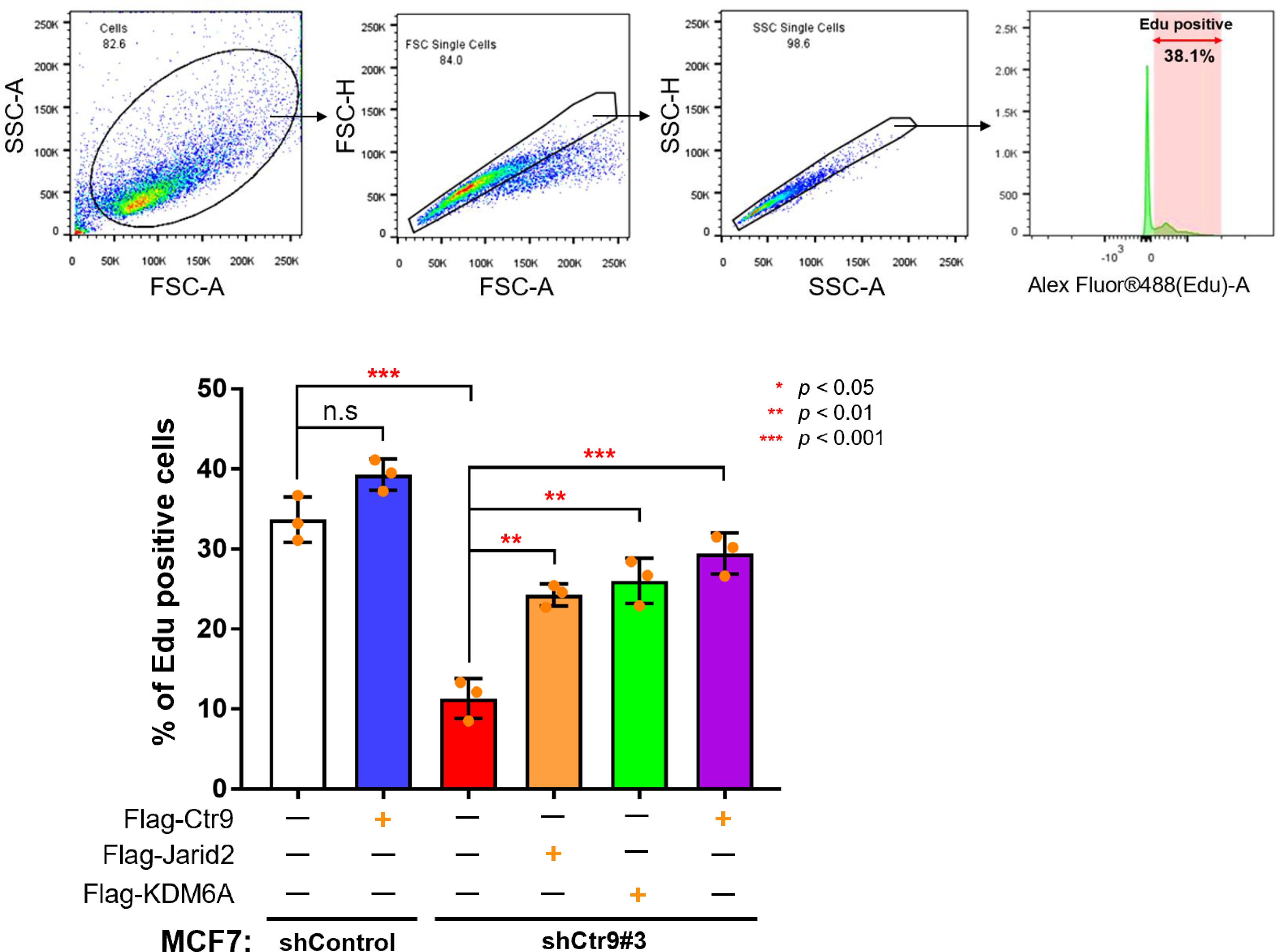
Exogenous expression of JARID2 or KDM6A attenuate the proliferation-defect of Ctr9 KD MCF7 cells. (Up) Gating strategy for Edu staining based flow cytometry analysis. (Bottom) Quantification of Edu positive cells in control KD or Ctr9 KD MCF7 cells transfected with blank vector or Flag-Jarid2/KDM6A/Ctr9 expressing plasmid. P-value of unpaired two tails t test with Welch correction were calculated.

**Figure S8 (related to Figure 7).**
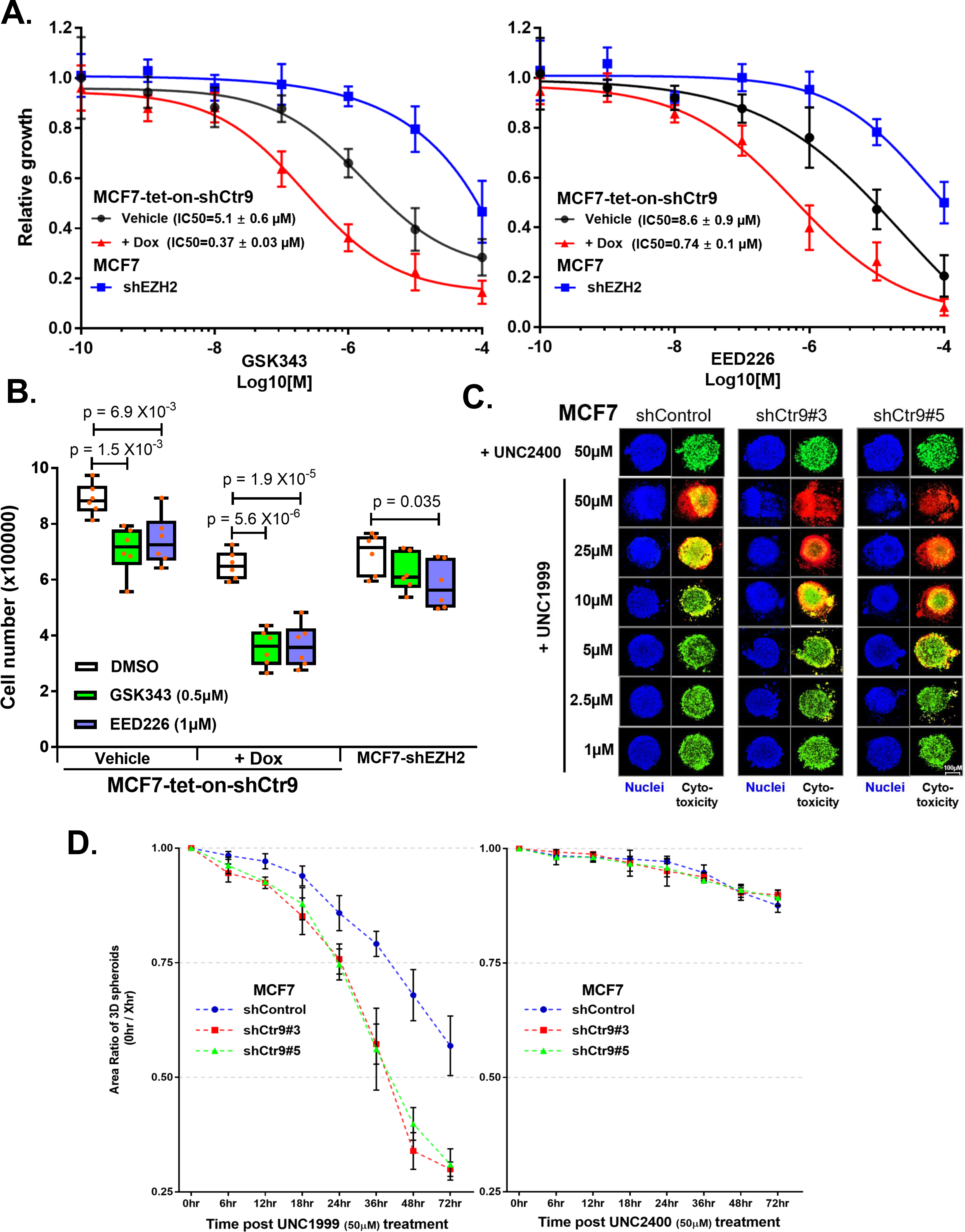
Depletion of CTR9 sensitizes MCF7 cells to two mechanistically distinct PRC2 inhibitors. A. Cell viability measured by MTT assays after treating MCF7-shEZH2 or MCF7-tet-on-shCtr9 cells (pretreated with vehicle or Dox) with increasing doses of SAM-based EZH2 inhibitor GSK343 (left) or PRC2 allosteric inhibitor EED226 (right). Data are represented as the mean±SD (n=3) normalized to 10^-10^ M treatment data. The IC_50_ was calculated using a non-linear regression model of log(inhibitor) response. B. Measurement of cell proliferation by cell counting with trypan blue exclusion after treating MCF7-shEZH2 or MCF7-tet-on-shCtr9 cells (pretreated with vehicle or Dox) with DMSO, 0.5 µM GSK343, or 1µM EED226 for four days. Data are represented as the mean ± SD (n=6). Adjusted p-value were calculated by multiple t test (unpaired, two tails) with Holm-Sidak correction. C. Representative confocal images of 3D spheroids (MCF7 shControl/ shCtr9#3/ shCtr9#5) after treated with ascending concentration of UNC1999 (EZH2i) for 2 days. UNC2400 (negative paralog) serves as a negative control. Nuclei were stained in blue. Live cells with ubiquitous esterase activity were in green; Dead cells with impaired cell membrane were in red. D. Time coursed measurement of 3D spheroids (MCF7-shControl/ shCtr9#3/ shCtr9#5) area ratio (Area _0hr_ / Area _Xhr_) after treated with 50 µM UNC1999 (EZH2i, left) or 50 µM UNC2400 (Negative paralog, right) for 0 – 72 hr. Data are represented as mean ± SD. (n=3)

**Table S1.**
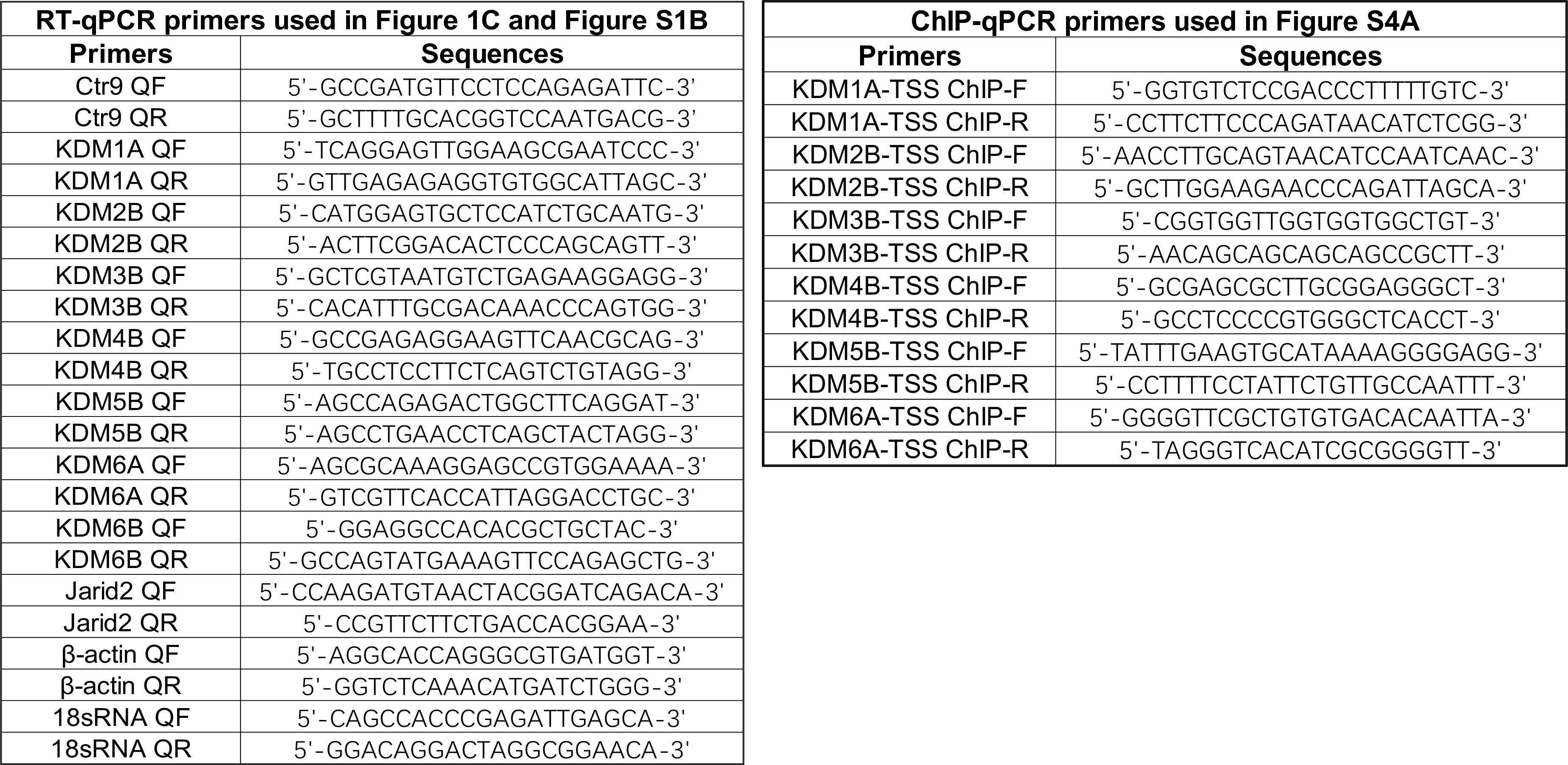
Sequence of RT-qPCR and ChIP-qPCR primers used in Figure 1C, Figure S1B and Figure S4B

## REFERENCES

1. Adam, F., Mark, D., Anne-Maud, F., Rory, J., Irwin, J., Paul, F., et.al., (2019) GENCODE reference annotation for the human and mouse genomes. Nucleic Acids Res. 47. D766– D773.

2. Agger, K., Cloos, P.A., Christensen, J., Pasini, D., Rose, S., Rappsilber, J., Issaeva, I., Canaani, E., Salcini, A.E., and Helin, K. (2007). UTX and JMJD3 are histone H3K27 demethylases involved in HOX gene regulation and development. Nature 449, 731–734.

3. Akanuma, T., Koshida, S., Kawamura, A., Kishimoto, Y., and Takada, S. (2007). Paf1 complex homologues are required for Notch-regulated transcription during somite segmentation. EMBO Rep. 8, 858–863.

4. Antonysamy, S., Condon, B., Druzina, Z., Bonanno, J.B., Gheyi, T., Zhang, F., MacEwan, I., Zhang, A., Ashok, S., Rodgers, L., et al. (2013). Structural context of disease-associated mutations and putative mechanism of autoinhibition revealed by X-ray crystallographic analysis of the EZH2-SET domain. PLoS One 8, e84147.

5. Ballare, C., Lange, M., Lapinaite, A., Martin, G.M., Morey, L., Pascual, G., Liefke, R., Simon, B., Shi, Y., Gozani, O., et al. (2012). Phf19 links methylated Lys36 of histone H3 to regulation of Polycomb activity. Nat. Struct. Mol. Biol. 19, 1257–1265.

6. Bamodu, O.A., Huang, W.C., Lee, W.H., Wu, A., Wang, L.S., Hsiao, M., Yeh, C.T., and Chao, T.Y. (2016). Aberrant KDM5B expression promotes aggressive breast cancer through MALAT1 overexpression and downregulation of hsa-miR-448. BMC Cancer 16, 160.

7. Ben, L., Cole, T., Mihai, P., Steven, L. S. (2009). Ultrafast and memory-efficient alignment of short DNA sequences to the human genome. BMC Genome Bio. 10, R25.

8. Brien, G.L., Gambero, G., O’Connell, D.J., Jerman, E., Turner, S.A., Egan, C.M., Dunne, E.J., Jurgens, M.C., Wynne, K., Piao, L., et al. (2012). Polycomb PHF19 binds H3K36me3 and recruits PRC2 and demethylase NO66 to embryonic stem cell genes during differentiation. Nat. Struct. Mol. Biol. 19, 1273–1281.

9. Cao, R., and Zhang, Y. (2004). SUZ12 is required for both the histone methyltransferase activity and the silencing function of the EED-EZH2 complex. Mol. Cell 15, 57–67.

10. Chaturvedi, D., Inaba, M., Scoggin, S., and Buszczak, M. (2016). Drosophila CG2469 Encodes a Homolog of Human CTR9 and Is Essential for Development. G3 (Bethesda) 6, 3849–3857.

11. Chen, S.L., Loffler, K.A., Chen, D., Stallcup, M.R., and Muscat, G.E. (2002). The coactivator-associated arginine methyltransferase is necessary for muscle differentiation: CARM1 coactivates myocyte enhancer factor-2. J. Biol. Chem. 277, 4324–4333.

12. Chu, X., Qin, X., Xu, H., Li, L., Wang, Z., Li, F., Xie, X., Zhou, H., Shen, Y., and Long, J. (2013). Structural insights into Paf1 complex assembly and histone binding. Nucleic Acids Res. 41, 10619–10629.

13. Ciferri, C., Lander, G.C., Maiolica, A., Herzog, F., Aebersold, R., and Nogales, E. (2012). Molecular architecture of human polycomb repressive complex 2. Elife 1, e00005.

14. Ciriello, G., Gatza, M.L., Beck, A.H., Wilkerson, M.D., Rhie, S.K., Pastore, A., Zhang, H., McLellan, M., Yau, C., Kandoth, C., et al. (2015). Comprehensive Molecular Portraits of Invasive Lobular Breast Cancer. Cell 163, 506–519.

15. Conway, E., Healy, E., and Bracken, A.P. (2015). PRC2 mediated H3K27 methylations in cellular identity and cancer. Curr. Opin. Cell Biol. 37, 42–48.

16. Conway, E., Jerman, E., Healy, E., Ito, S., Holoch, D., Oliviero, G., Deevy, O., Glancy, E., Fitzpatrick, D.J., Mucha, M., et al. (2018). A Family of Vertebrate-Specific Polycombs Encoded by the LCOR/LCORL Genes Balance PRC2 Subtype Activities. Mol. Cell 70, 408–421.

17. Ezponda, T., Dupere-Richer, D., Will, C.M., Small, E.C., Varghese, N., Patel, T., Nabet, B., Popovic, R., Oyer, J., Bulic, M., et al. (2017). UTX/KDM6A Loss Enhances the Malignant Phenotype of Multiple Myeloma and Sensitizes Cells to EZH2 inhibition. Cell Rep. 21, 628–640.

18. Gong, Y., Huo, L., Liu, P., Sneige, N., Sun, X., Ueno, N.T., Lucci, A., Buchholz, T.A., Valero, V., and Cristofanilli, M. (2011). Polycomb group protein EZH2 is frequently expressed in inflammatory breast cancer and is predictive of worse clinical outcome. Cancer 117, 5476–5484.

19. Grijzenhout, A., Godwin, J., Koseki, H., Gdula, M.R., Szumska, D., McGouran, J.F., Bhattacharya, S., Kessler, B.M., Brockdorff, N., and Cooper, S. (2016). Functional analysis of AEBP2, a PRC2 Polycomb protein, reveals a Trithorax phenotype in embryonic development and in ESCs. Development 143, 2716–2723.

20. Grosselin, K., Durand, A., Marsolier, J., Poitou, A., Marangoni, E., Nemati, F., Dahmani, A., Lameiras, S., Reyal, F., Frenoy, O., et al. (2019). High-throughput single-cell ChIP-seq identifies heterogeneity of chromatin states in breast cancer. Nat. Genet. 51, 1060–1066.

21. Gu, Z., Roland, E., Matthias, S. (2016) Complex heatmaps reveal patterns and correlations in multidimensional genomic data. Bioinformatics 32, 2847–2849.

22. Hanks, S., Perdeaux, E.R., Seal, S., Ruark, E., Mahamdallie, S.S., Murray, A., Ramsay, E., Del Vecchio Duarte, S., Zachariou, A., de Souza, B., et al. (2014). Germline mutations in the PAF1 complex gene CTR9 predispose to Wilms tumour. Nat. Commun. 5, 4398.

23. Hirukawa, A., Smith, H.W., Zuo, D., Dufour, C.R., Savage, P., Bertos, N., Johnson, R.M., Bui, T., Bourque, G., Basik, M., et al. (2018). Targeting EZH2 reactivates a breast cancer subtype-specific anti-metastatic transcriptional program. Nat. Commun. 9, 2547.

24. Holoch, D., and Margueron, R. (2017). Mechanisms Regulating PRC2 Recruitment and Enzymatic Activity. Trends Biochem. Sci. 42, 531–542.

25. Hosogane, M., Funayama, R., Shirota, M., and Nakayama, K. (2016). Lack of Transcription Triggers H3K27me3 Accumulation in the Gene Body. Cell Rep. 16, 696–706.

26. Jaehning, J.A. (2010). The Paf1 complex: platform or player in RNA polymerase II transcription? Biochim Biophys. Acta. 1799, 379–388.

27. Jake, R. C., Alexander, L., Nils, G. (2017). UpSetR: an R package for the visualization of intersecting sets and their properties. Bioinformatics 33, 2938–2940,

28. Kasinath, V., Faini, M., Poepsel, S., Reif, D., Feng, X.A., Stjepanovic, G., Aebersold, R., and Nogales, E. (2018). Structures of human PRC2 with its cofactors AEBP2 and JARID2. Science 359, 940–944.

29. Kim, K.H., and Roberts, C.W. (2016). Targeting EZH2 in cancer. Nat. Med. 22, 128–134.

30. Kleer, C.G., Cao, Q., Varambally, S., Shen, R., Ota, I., Tomlins, S.A., Ghosh, D., Sewalt, R.G., Otte, A.P., Hayes, D.F., et al. (2003). EZH2 is a marker of aggressive breast cancer and promotes neoplastic transformation of breast epithelial cells. Proc. Natl. Acad. Sci. U S A 100, 11606–11611.

31. Konze, K.D., Ma, A., Li, F., Barsyte-Lovejoy, D., Parton, T., Macnevin, C.J., Liu, F., Gao, C., Huang, X.P., Kuznetsova, E., et al. (2013). An orally bioavailable chemical probe of the Lysine Methyltransferases EZH2 and EZH1. ACS Chem. Biol. 8, 1324–1334.

32. Kooistra, S.M., and Helin, K. (2012). Molecular mechanisms and potential functions of histone demethylases. Nat Rev Mol. Cell Biol. 13, 297–311.

33. Lan, F., Nottke, A.C., and Shi, Y. (2008). Mechanisms involved in the regulation of histone lysine demethylases. Curr. Opin. Cell Biol. 20, 316–325.

34. Laugesen, A., Hojfeldt, J.W., and Helin, K. (2019). Molecular Mechanisms Directing PRC2 Recruitment and H3K27 Methylation. Mol. Cell 74, 8–18.

35. Malte, B., Paola, P., Valerio, D. C., Enrique, B., Paul, C., Pedro, V., Arantxa, G., Sergi, A., Bernhard, P., Michael, W., Luciano, D. C. (2016). EPOP Functionally Links Elongin and Polycomb in Pluripotent Stem Cells. Mol. Cell 64, 645–358.

36. Maes, T., et al., KDM1 histone lysine demethylases as targets for treatments of oncological and neurodegenerative disease. Epigenomics, 2015. 7(4): p. 609–26.

37. Massoni-Laporte, A., Perrot, M., Ponger, L., Boucherie, H., and Guieysse-Peugeot, A.L. (2012). Proteome analysis of a CTR9 deficient yeast strain suggests that Ctr9 has function(s) independent of the Paf1 complex. Biochim. Biophys. Acta. 1824, 759–768.

38. Michael, H. K., Jamie, J. N., Steve, B., Ye, Z., David, A. O., Nynke, L. van B., Richard, A., Y., et al., (2010). Mediator and cohesin connect gene expression and chromatin architecture. Nature 467, 430–435.

39. Michael, L., Robert, G., Vincent, C. (2009). rtracklayer: an R package for interfacing with genome browsers. Bioinformatics 25, 1841–1842.

40. Michael, L., Wolfgang, H., Hervé, P., Patrick, A., Marc, C., Robert, G., Martin, T. M., Vincent, C.(2013).Software for Computing and Annotating Genomic Ranges. PLoS Comput. Biol. 9, e1003118.

41. Moore, H.M., Gonzalez, M.E., Toy, K.A., Cimino-Mathews, A., Argani, P., and Kleer, C.G. (2013). EZH2 inhibition decreases p38 signaling and suppresses breast cancer motility and metastasis. Breast Cancer Res. Treat 138, 741–752.

42. Myriam, H., Ruth, K., Robert, J. F., Paul, G., Nathaniel, H. (2014). Cell type–specific mRNA purification by translating ribosome affinity purification (TRAP). Nature Protocols 9, 1282–1291.

43. Nowak, R.P., Tumber, A., Johansson, C., Che, K.H., Brennan, P., Owen, D., and Oppermann, U. (2016). Advances and challenges in understanding histone demethylase biology. Curr. Opin. Chem. Biol. 33, 151–159.

44. Ntziachristos, P., Tsirigos, A., Welstead, G.G., Trimarchi, T., Bakogianni, S., Xu, L., Loizou, E., Holmfeldt, L., Strikoudis, A., King, B., et al. (2014). Contrasting roles of histone 3 lysine 27 demethylases in acute lymphoblastic leukaemia. Nature 514, 513–517.

45. Oksuz, O., Narendra, V., Lee, C.H., Descostes, N., LeRoy, G., Raviram, R., Blumenberg, L., Karch, K., Rocha, P.P., Garcia, B.A., et al. (2018). Capturing the Onset of PRC2-Mediated Repressive Domain Formation. Mol. Cell 70, 1149–1162.

46. Pasini, D., Bracken, A.P., Jensen, M.R., Lazzerini Denchi, E., and Helin, K. (2004). Suz12 is essential for mouse development and for EZH2 histone methyltransferase activity. EMBO J. 23, 4061–4071.

47. Pasini, D., Cloos, P.A., Walfridsson, J., Olsson, L., Bukowski, J.P., Johansen, J.V., Bak, M., Tommerup, N., Rappsilber, J., and Helin, K. (2010). JARID2 regulates binding of the Polycomb repressive complex 2 to target genes in ES cells. Nature 464, 306–310.

48. Peng, J.C., Valouev, A., Swigut, T., Zhang, J., Zhao, Y., Sidow, A., and Wysocka, J. (2009). Jarid2/Jumonji coordinates control of PRC2 enzymatic activity and target gene occupancy in pluripotent cells. Cell 139, 1290–1302.

49. Qi, W., Zhao, K., Gu, J., Huang, Y., Wang, Y., Zhang, H., Zhang, M., Zhang, J., Yu, Z., Li, L., et al. (2017). An allosteric PRC2 inhibitor targeting the H3K27me3 binding pocket of EED. Nat. Chem. Biol. 13, 381–388.

50. Riising, E.M., Comet, I., Leblanc, B., Wu, X., Johansen, J.V., and Helin, K. (2014). Gene silencing triggers polycomb repressive complex 2 recruitment to CpG islands genome wide. Mol. Cell 55, 347–360.

51. Sanulli, S., Justin, N., Teissandier, A., Ancelin, K., Portoso, M., Caron, M., Michaud, A., Lombard, B., da Rocha, S.T., Offer, J., et al. (2015). Jarid2 Methylation via the PRC2 Complex Regulates H3K27me3 Deposition during Cell Differentiation. Mol. Cell 57, 769–783.

52. Schmitges, F.W., Prusty, A.B., Faty, M., Stutzer, A., Lingaraju, G.M., Aiwazian, J., Sack, R., Hess, D., Li, L., Zhou, S., et al. (2011). Histone methylation by PRC2 is inhibited by active chromatin marks. Mol. Cell 42, 330–341.

53. Shen, X., Kim, W., Fujiwara, Y., Simon, M.D., Liu, Y., Mysliwiec, M.R., Yuan, G.C., Lee, Y., and Orkin, S.H. (2009). Jumonji modulates polycomb activity and self-renewal versus differentiation of stem cells. Cell 139, 1303–1314.

54. Shpargel, K.B., Starmer, J., Yee, D., Pohlers, M., and Magnuson, T. (2014). KDM6 demethylase independent loss of histone H3 lysine 27 trimethylation during early embryonic development. PLoS Genet. 10, e1004507.

55. Smits, A.H., Jansen, P.W., Poser, I., Hyman, A.A., and Vermeulen, M. (2013). Stoichiometry of chromatin-associated protein complexes revealed by label-free quantitative mass spectrometry-based proteomics. Nucleic Acids Res. 41, e28.

56. Streubel, G., Watson, A., Jammula, S.G., Scelfo, A., Fitzpatrick, D.J., Oliviero, G., McCole, R., Conway, E., Glancy, E., Negri, G.L., et al. (2018). The H3K36me2 Methyltransferase Nsd1 Demarcates PRC2-Mediated H3K27me2 and H3K27me3 Domains in Embryonic Stem Cells. Mol. Cell 70, 371–379.

57. Taube, J.H., Sphyris, N., Johnson, K.S., Reisenauer, K.N., Nesbit, T.A., Joseph, R., Vijay, G.V., Sarkar, T.R., Bhangre, N.A., Song, J.J., et al. (2017). The H3K27me3-demethylase KDM6A is suppressed in breast cancer stem-like cells, and enables the resolution of bivalency during the mesenchymal-epithelial transition. Oncotarget 8, 65548–65565.

58. Tomson, B.N., and Arndt, K.M. (2013). The many roles of the conserved eukaryotic Paf1 complex in regulating transcription, histone modifications, and disease states. Biochim. Biophys. Acta. 1829, 116–126.

59. Van der Meulen, J., Sanghvi, V., Mavrakis, K., Durinck, K., Fang, F., Matthijssens, F., Rondou, P., Rosen, M., Pieters, T., Vandenberghe, P., et al. (2015). The H3K27me3 demethylase UTX is a gender-specific tumor suppressor in T-cell acute lymphoblastic leukemia. Blood 125, 13–21.

60. Van Mierlo, G., Veenstra, G.J.C., Vermeulen, M., and Marks, H. (2019). The Complexity of PRC2 Subcomplexes. Trends Cell. Biol. (Cell Reviews) 29, 660–671.

61. Van Oss, S.B., Cucinotta, C.E., and Arndt, K.M. (2017). Emerging Insights into the Roles of the Paf1 Complex in Gene Regulation. Trends Biochem. Sci. 42, 788–798.

62. Verma, S.K., Tian, X., LaFrance, L.V., Duquenne, C., Suarez, D.P., Newlander, K.A., Romeril, S.P., Burgess, J.L., Grant, S.W., Brackley, J.A., et al. (2012). Identification of Potent, Selective, Cell-Active Inhibitors of the Histone Lysine Methyltransferase EZH2. ACS Med. Chem. Lett. 3, 1091–1096.

63. Vos, S.M., Farnung, L., Boehning, M., Wigge, C., Linden, A., Urlaub, H., and Cramer, P. (2018). Structure of activated transcription complex Pol II-DSIF-PAF-SPT6. Nature 560, 607–612.

64. Wickham, H. (2016). ggplot2: Elegant Graphics for Data Analysis. Springer-Verlag: New York, USA.

65. Wu, H., Zeng, H., Dong, A., Li, F., He, H., Senisterra, G., Seitova, A., Duan, S., Brown, P.J., Vedadi, M., et al. (2013). Structure of the catalytic domain of EZH2 reveals conformational plasticity in cofactor and substrate binding sites and explains oncogenic mutations. PLoS One 8, e83737.

66. Wu, J., and Xu, W. (2012) Histone H3R17me2a mark recruits human RNA Polymerase Associated Factor 1 Complex to activate transcription. PNAS 109, 5675–5680.

67. Xie, Y., Zheng, M., Chu, X., Chen, Y., Xu, H., Wang, J., Zhou, H., and Long, J. (2018). Paf1 and Ctr9 subcomplex formation is essential for Paf1 complex assembly and functional regulation. Nat. Commun. 9, 3795.

68. Xu, B., Konze, K.D., Jin, J., and Wang, G.G. (2015). Targeting EZH2 and PRC2 dependence as novel anticancer therapy. Exp. Hematol 43, 698–712.

69. Yu, J.R., Lee, C.H., Oksuz, O., Stafford, J.M., and Reinberg, D. (2019). PRC2 is high maintenance. Genes Dev. (Reviews) 33, 1–33.

70. Yu, M., Yang, W., Ni, T., Tang, Z., Nakadai, T., Zhu, J., and Roeder, R.G. (2015). RNA polymerase II-associated factor 1 regulates the release and phosphorylation of paused RNA polymerase II. Science 350, 1383–1386.

71. Yu, Y., Qi, J., Xiong, J., Jiang, L., Cui, D., He, J., Chen, P., Li, L., Wu, C., Ma, T., et al. (2019b). Epigenetic Co-Deregulation of EZH2/TET1 is a Senescence-Countering, Actionable Vulnerability in Triple-Negative Breast Cancer. Theranostics 9, 761–777.

72. Zeng, H., Lu, L., Chan, N.T., Horswill, M., Ahlquist, P., Zhong, X., and Xu, W. (2016). Systematic identification of Ctr9 regulome in ERalpha-positive breast cancer. BMC Genomics 17, 902.

73. Zeng, H., and Xu, W. (2015). Ctr9, a key subunit of PAFc, affects global estrogen signaling and drives ERalpha-positive breast tumorigenesis. Genes Dev. 29, 2153–2167.

74. Zeng, H., and Xu, W. (2016). Gene expression profiling of Ctr9-regulated transcriptome in ERalpha-positive breast cancer. Genom. Data 7, 103–104.

75. Zhang, K., Haversat, J.M., and Mager, J. (2013). CTR9/PAF1c regulates molecular lineage identity, histone H3K36 trimethylation and genomic imprinting during preimplantation development. Dev. Biol. 383, 15–27.

76. Zhang, Y., Liu, T., Clifford, A. M., Jérôme, E., David, S. J., Bradley, E. B., Chad, N., Richard, M. M., Myles, B., Li, W., Liu, X. S. (2008) Model-based Analysis of ChIP-Seq (MACS). BMC Genome Bio. 9, R137.

77. Zhu, B., Mandal, S.S., Pham, A.D., Zheng, Y., Erdjument-Bromage, H., Batra, S.K., Tempst, P., and Reinberg, D. (2005). The human PAF complex coordinates transcription with events downstream of RNA synthesis. Genes Dev. 19, 1668–1673.

